# Location specific small RNA annealing to the HCV 5’ UTR promotes Hepatitis C Virus replication by favoring IRES formation and stimulating virus translation

**DOI:** 10.1101/2020.03.25.008417

**Authors:** Rasika D. Kunden, Sarah Ghezelbash, Juveriya Q. Khan, Joyce A. Wilson

## Abstract

Hepatitis C virus (HCV) genome replication requires annealing of a liver specific small-RNA, miR-122 to 2 sites on 5’ untranslated region (UTR). Annealing has been reported to a) stabilize the genome, b) promote translation, and c) induce the canonical HCV 5’ UTR Internal Ribosome Entry Site (IRES) structure. In this report we identify the relative impact of small RNA annealing on the three functions ascribed to miR-122 and generate a mechanistic model for miR-122 promotion of HCV. First, we identified that perfectly complementary small RNAs that anneal to different locations on the HCV 5’ UTR stimulate replication with varying efficiencies and mapped the region on the HCV genome to which small RNA annealing promotes virus replication. Second, by using a panel of small RNAs that promote with varying efficiencies we link HCV replication induction with translation stimulation and 5’ UTR RNA structure modifications. However, replication promotion was not linked to genome stabilization since all small RNAs tested could stabilize the viral genome regardless of their ability to promote replication. Thus, we propose that miR-122 annealing promotes HCV replication primarily by activating the HCV IRES and stimulating translation, and that miR-122-induced HCV genome stabilization is insufficient alone but enhances virus replication.

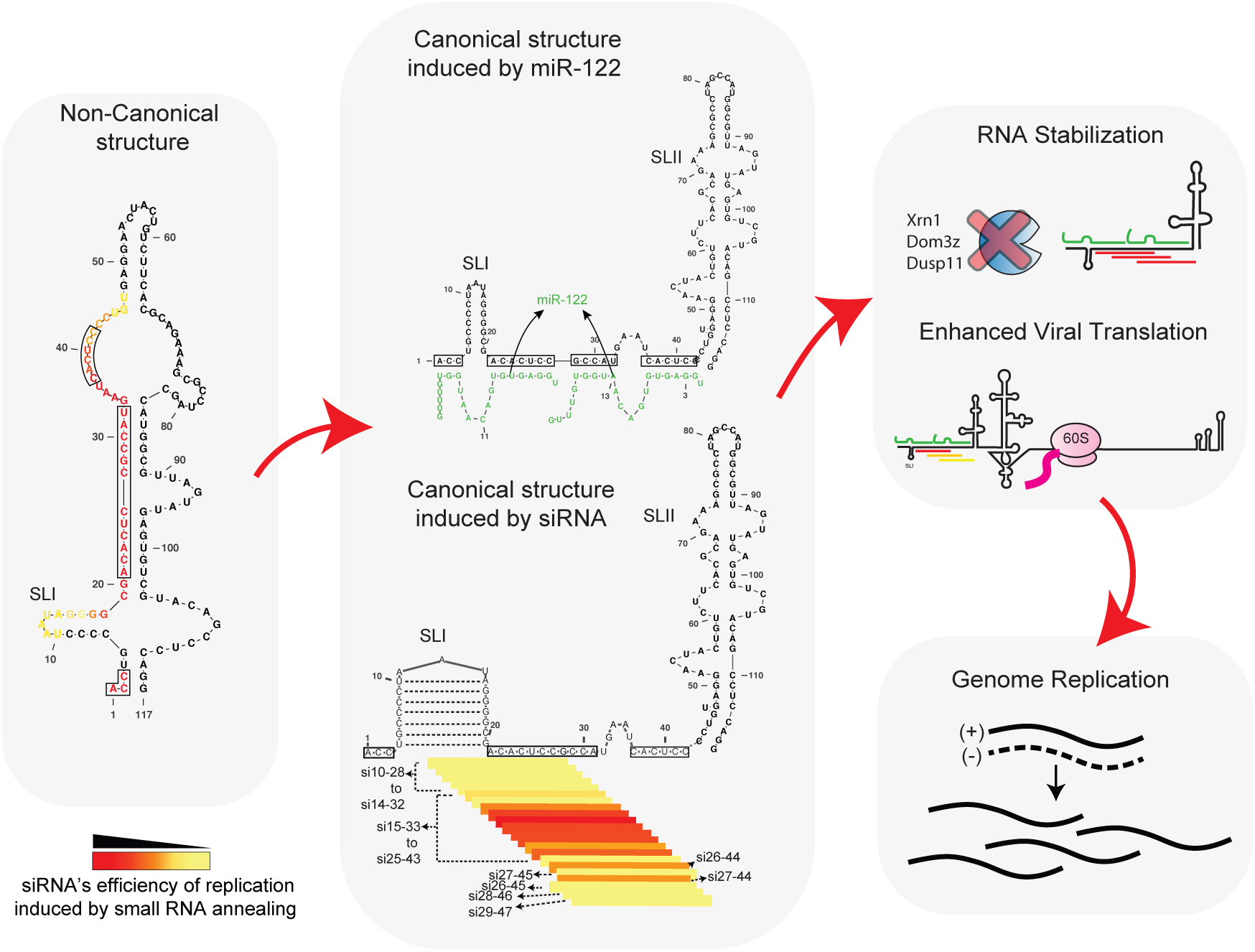
Graphical Abstract.

## INTRODUCTION

Hepatitis C Virus (HCV) is a flavivirus that causes chronic infections of the liver and can lead to liver cirrhosis and hepatocellular carcinoma (1, 2). The genome of HCV is a 9.6 kb long positive sense RNA that consists of a 5’ untranslated region (UTR), a polyprotein coding region, and a 3’UTR region. The 5’ and 3’ UTRs are highly structured and required for genome translation and replication (3, 4).

The 5’UTR is a structured RNA that forms 4 stem loops (SL), SLI, SLII, SLIII and SLIV. SLII, SLIII and SLIV comprise the internal ribosomal entry site (IRES) that drives cap-independent HCV translation (Figure 1A) (5, 6). SLII is divided into two parts, SLIIa which induces SLII to form a bent structure that directs SLIIb, to the ribosomal E-site in the head region of the 40S subunit, facilitating 80S ribosome assembly (7–9). The first 42 nucleotides on the 5’UTR are not considered part of the IRES and forms SLI and an RNA structured element created by annealing of two copies of microRNA-122 (miR-122), a host microRNA found in human liver cells (10–13). This tri-molecular structure is required for virus replication (11–14).

**Figure 1:**
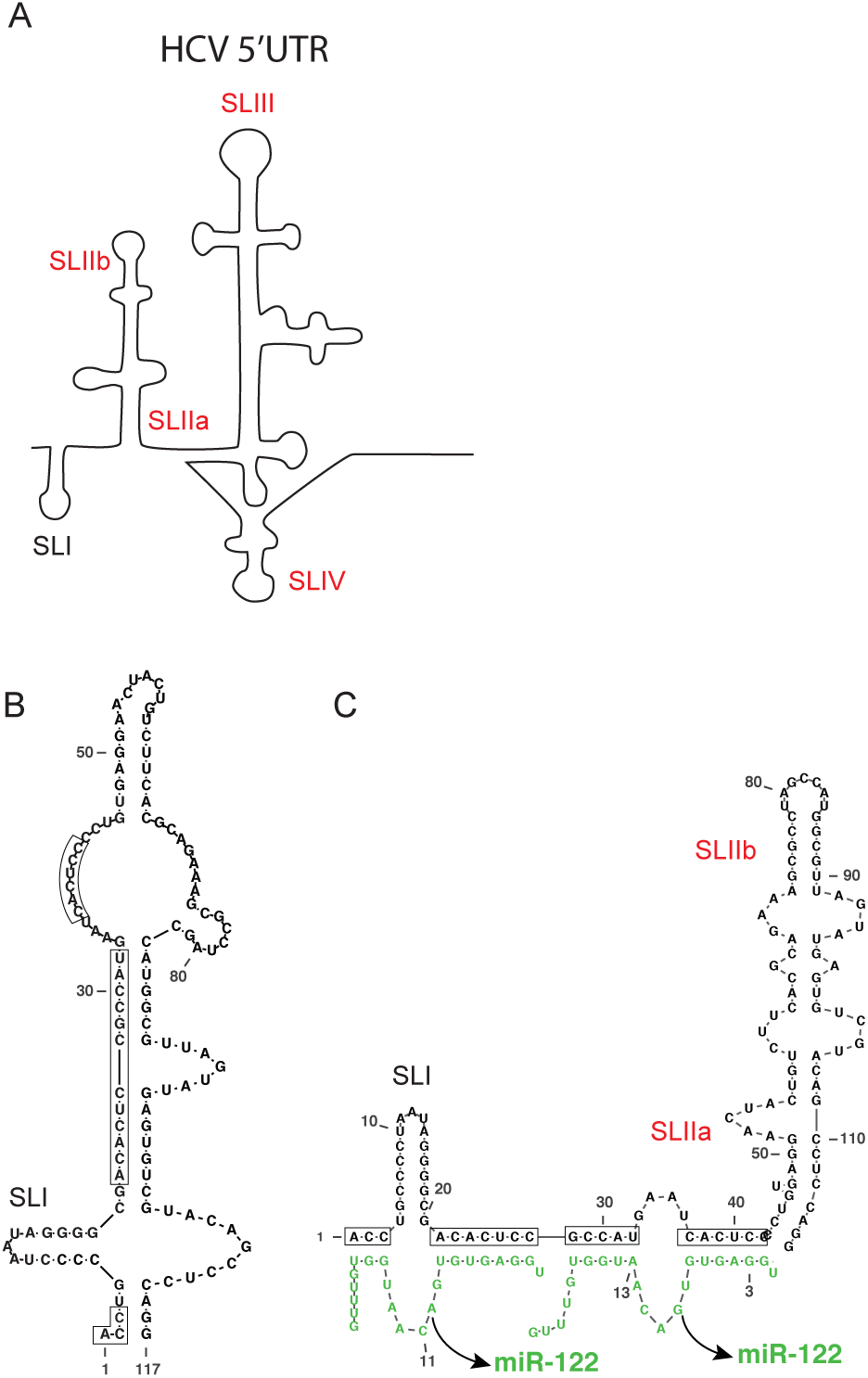
HCV 5’ UTR RNA structures. (A). Schematic representation of HCV 5’UTR stem loops; SLI, SLIIa, SLIIb, SLIII and SLIV are indicated. Predicted structure of the 5’ 117 nucleotide RNA fragment alone forms a non-canonical RNA structure (B). and after annealing of 2 copies of miR-122 forms the canonical structure (C). The miR-122 binding nucleotides are shown within black boxes. Stem loops indicated in Red are parts of IRES.

MicroRNAs (miRNAs) are small RNAs about 21-23 nucleotides long and are central to mRNA regulated by miRNA gene silencing (15, 16). miRNAs silence genes in association with a host Argonaute (Ago) protein within an RNA induced silencing complex (RISC) and directs the protein complex to the 3’ UTR of an mRNA by annealing with imperfect sequence complementarity (17). miRNAs target an mRNA by annealing to an approximately 6-7 nucleotide seed site binding at the 5’ end of the miRNA and an accessory site binding on the 3’end (18). This leaves a loop of mismatched nucleotides between the seed and accessory sites and overhangs on the 5’ and 3’ ends of the miRNA that do not bind to the target mRNA. Such complex miRNA:target RNA interactions regulates the expression of multiple mRNAs by inducing translation suppression and degradation of target mRNA (19). Two copies of miR-122 anneal in conjunction with Ago to the HCV genome in a similar manner, including seed and accessory binding sites (20) but in this case promotes instead of silencing the viral genome.

HCV replication is undetectable in the absence of miR-122 and the mechanism behind miR-122’s stimulatory effect is not fully understood. miR-122 stabilizes the viral RNA by protecting it from degradation by host exonuclease Xrn1, and phosphatases Dom3Z and DUSP11 (21–23). However, simultaneous knock-down of these three enzymes cannot completely rescue HCV replication in the absence of miR-122, suggesting that other roles exist (23). miR-122 also promotes HCV translation and recent reports hypothesize that miR-122 modifies the HCV genome to induce the canonical 5’UTR IRES structure (Figure 1B and C) (12, 13, 24). Finally, miR-122 has been reported to directly induce genome amplification (25, 26). However, the relative impact of these functions on miR-122 directed HCV replication promotion is unknown (27).

We previously showed that annealing of small interfering RNAs (siRNA) to the HCV 5’ UTR can mimic the pro-viral activity of miR-122 (13). Like miRNAs, siRNAs are also 21–23 nucleotides in length and associate with Ago proteins, but based on associating with Ago2 and perfect sequence match with their targets, induce mRNA cleavage and gene knockdown (28, 29). However, when siRNA cleavage activity was blocked by using Ago2 knockout cells, siRNAs that anneal to the miR-122 binding region on the HCV genome they promoted virus replication, some as efficiently as miR-122. That siRNA annealing promoted HCV, suggested that the complex annealing pattern formed by miR-122 on the HCV genome was not required for the pro-viral activity. It also provided a method to assess the impact of small RNA annealing to other locations on the genome on HCV replication (13). Using a panel of HCV genome-targeting siRNAs, we found that annealing between nucleotides 1 and 44 in HCV 5’UTR, promoted HCV replication, and annealing within the IRES, NS5B and 3’UTR regions did not. We also found that siRNAs that annealed to different locations on the 5’ UTR promoted virus replication with different efficiencies and the efficiency correlated with their ability to stimulate translation and their predicted ability to induce the canonical SLII structure of the HCV IRES. Finally, like miR-122, annealing of the siRNAs to the 5’ UTR also stabilized the viral genome, but the siRNAs did so regardless of whether they could promote replication, suggesting that genome stabilization alone is insufficient for HCV replication promotion. Thus, our current model for the pro-viral mechanism of miR-122 replication posits that its primary role is to stimulate translation by inducing a 5’ UTR RNA structural switch element to form the canonical HCV IRES structure, and that genome stabilization is its secondary role that is insufficient alone, but enhances replication induced by translation stimulation.

## MATERIALS AND METHODS

### Plasmids

Full-length HCV Renilla Luciferase (Rluc) reporter genome constructs pJ6/JFH-1 RLuc (p7-RLuc2A) and the non-replicative version pJ6/JFH-1 RLuc (p7-RLuc2A) GNN were provided by Dr. C. M. Rice (30). For translation suppression assays the miR-122 suppression firefly luciferase (Fluc) reporter plasmid, pFLuc JFH-1 5’UTR × 2 (11) was modified to contain a single copy of the complete 5’UTR (pFluc JFH-1 5’UTR) or NS5B-3’UTR (pFluc JFH-1 NS5B-3’UTR) region of the genome by replacing the fragment between restriction sites SpeI and SacII. To generate the complete 5’ UTR fragment we used PCR and the forward primer 5’GCCACTAGTACGACGGCCAGTGAATTC3’; and reverse primer 5’CAGCCGCGGATCGATGACCTTACCCACG3’, to generate the NS5B-3’UTR region we used forward primer 5’GCCACTAGTAATGTGTCTGTGGCGTTGG3’; reverse primer 5’CAGCCGCGGAAACAGCTATGACCATGA3’ and the pJ6/JFH-1 RLuc (p7-RLuc2A) plasmid as a template. Each Forward primer has sequence for restriction site, SpeI, and each Reverse primer has sequence for SacII. The control plasmid pRL-TK was obtained from Promega (Madison, USA). A pT7 Fluc containing plasmid, herein called pT7 Fluc (Promega, Madison, USA), was used to generate control mRNA for translation assays.

### In vitro RNA transcription

To generate full length viral RNA the plasmid pJ6/JFH-1 RLuc (p7-RLuc2A) or related mutants were linearized by digestion with XbaI and RNA was made by using the MEGA Script T7 High Yield Transcription Kit (Life Technologies, Burlington, Canada). The transcription process was performed using the suggested manufacturer’s protocol. Fluc mRNA transcript was prepared by digesting the plasmid, pT7 Fluc mRNA, with XmnI and mRNA was prepared using the mMessage mMachine mRNA synthesis kit (Life Technologies, Burlington, Canada) using manufacturer’s protocol.

### Small interfering RNA (siRNA) design and sequence

HCV targeting siRNAs that anneal to regions in IRES, NS5B coding region and 3’UTR were designed using online software, i-score, https://www.med.nagoya-u.ac.jp/neurogenetics/i_Score/i_score.html (31). The sequence of siRNA JFH-1 6367 (si6367) was adapted from the siRNA described previously to inhibit the HCV con1 genotype, by modifying the sequence to match the same region in JFH-1 GACCCACAAACACCAAUUCCC (32). The control siRNA (siControl) target sequence is GAGAGUCAGUCAGCUAAUCA and does not anneal to the virus genome. All siRNAs that anneal to the HCV genome were designed to have 21 nucleotides; 19 nucleotides complementary to target site and 2 UU overhangs on the 3’ end for incorporating into RISC complex, unless stated otherwise (Supplementary Table 3). These siRNAs were synthesized by GE Lifesciences Dharmacon. Anti-miR-122, miRIDIAN microRNA Human hsa-miR-122-5p-Hairpin Inhibitor (IH-300591-06-0050), were purchased from Dharmacon Horizon Discoveries (Chicago, USA).

### Cell Culture

miR-122 knockout (miR-122 KO) Huh-7.5 (33), Ago2 knockout (Ago2 KO) Huh-7.5 cells (13) and DROSHA/Ago2 KO cells were cultured in Dulbecco’s modified Eagle medium (DMEM) supplemented with 10% fetal bovine serum, 0.1 nM non-essential amino acids (Wisent, Montreal, Canada) and 100 ug/ml Pen/Strep (Invitrogen, Burlingtion, Canada). miR-122 knockout Huh-7.5 cells were a kind gift from Dr Matthew Evans. DROSHA/Ago2 KO Huh 7.5 cells were generated from DROSHA KO Huh 7.5 cells (34) (a gift from Dr. Charlie Rice) by using the CRISPR-Cas9 genome editing techniques (35).

### siRNA suppression assay

To assess the ability of an siRNA to suppress mRNA translation and assess whether the siRNAs are functional in the RISC, we assessed their impact in a transient suppression assay (Figure 2B). For this assay we used reporter plasmids (pFluc JFH-1 5’UTR **/** pFuc JFH-1 NS5B-3’UTR) that express Fluc mRNAs containing the siRNA target sequence from HCV in their 3’UTRs. We also used an Rluc expressing control plasmid, pRL-TK to normalize transfection efficiency (Promega, Madison, USA). The day before transfection 8 x 10^4^ miR-122 KO cells/well were plated in a 24 well dish and incubated overnight. The next day, the cells were transfected with 100ng of each of pRL-TK and pFluc JFH-1 (5’UTR or NS5B-3’UTR) and 0.1pmol of a particular test siRNA. The transfection mixture was prepared using 1 μl lipofectamine 2000 according to the suggested manufacture’s protocol (Life Technologies, Burlington, Canada). The cells were incubated at 37°C, 5% CO2 after transfections and after 48 hours were lysed using passive lysis buffer and assayed for Fluc and Rluc activity using a dual luciferase assay kit (Promega, Madison, USA) (13).

**Figure 2:**
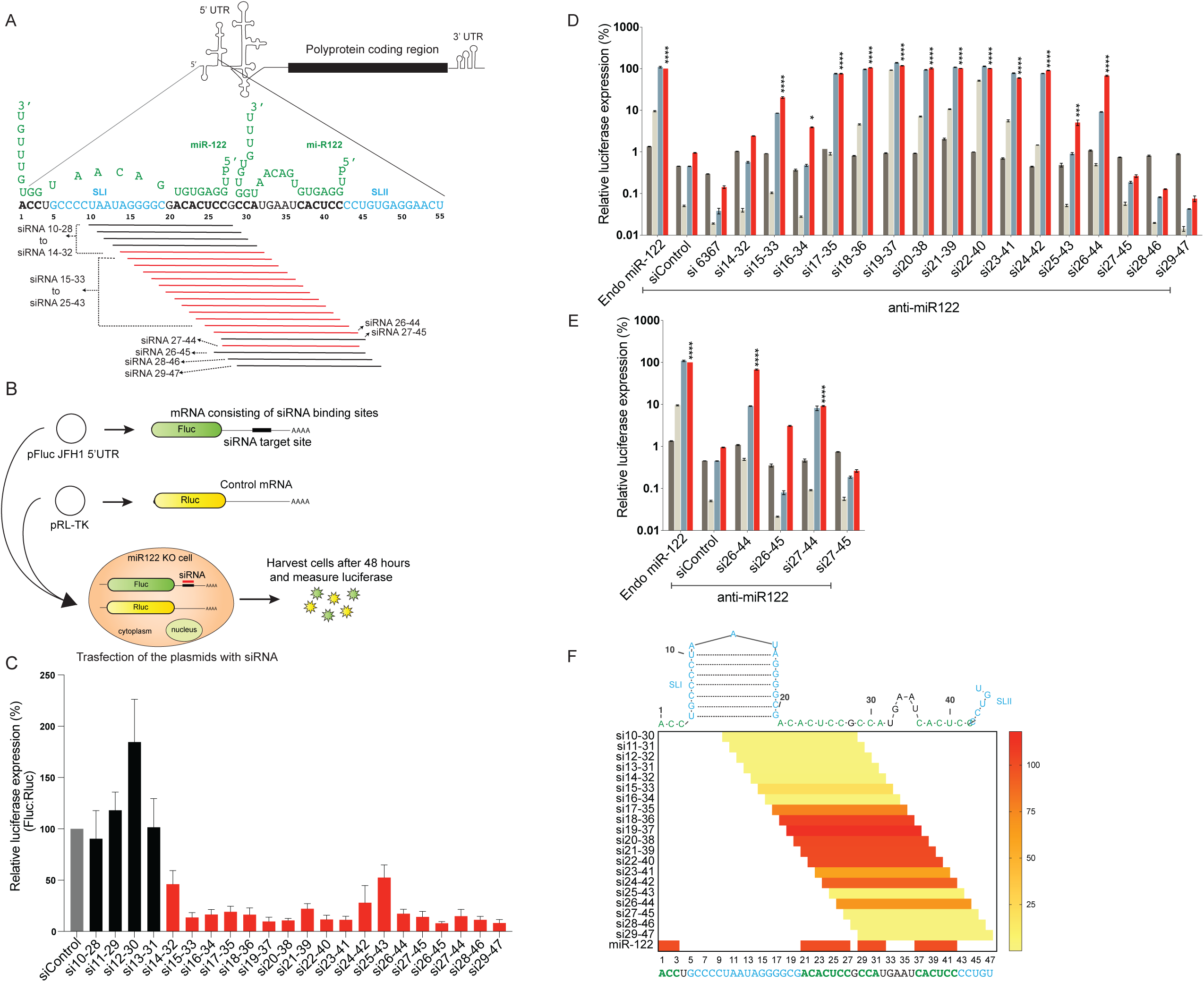
HCV replication promotion by siRNAs binding to nucleotides 10-47 in the 5’UTR. (A) Diagrammatic representation of HCV genome showing 5’UTR polyprotein coding region and 3’ UTR. The first 55 nucleotides of HCV 5’ UTR are shown interacting with 2 copies of miR-122 (green). siRNAs designed to walk the 5’UTR with single nucleotide resolution are represented. Black lines represent siRNAs that do not promote replication and red lines represent ones that do. (B) Diagram depicting the siRNA suppression assay. (C) siRNA suppression assay results with siControl (grey bar), bars are colour coded based on their suppression activities. siRNAs that do not suppress translation (black bars) and that suppress translation (red bars). (D and E) The pro-viral activity of the siRNAs was assessed using HCV replication assays in which the activity of miR-122 is antagonized using anti-miR-122. Ago2 knockout cells were co-electroporated with HCV J6/ JFH-1(p7-Rluc2a) RNA, anti-miR-122 and the indicated siRNA and harvested at 2 hours (grey bars), 24 hours (light grey bars), 48 hours (blue bars) and 72 hours (red bars) post-electroporation. HCV replication was measured based on Rluc expression and is presented as % relative to Rluc expression from HCV RNA supported by endogenous miR-122 at 72 hours post electroporation (Endo miR-122). The data are the average of at least 3 independent experiments and error bars represent the standard deviation. Statistical significance was determine using one-way ANOVA on the relative 72-hour values where, *P<0.0332; **P< 0.0021; ***P < 0.0002; ****P < 0.0001. (F). Heat map showing small-RNA efficiencies to promote virus replication. Red signal means highest efficiency and yellow signals mean lowest efficiencies. Secondary structure of HCV 1-47 nucleotides is shown to visualize the effect of small-RNA location binding on HCV replication efficacy.

### HCV replication Assay

Ago2 KO Huh-7.5 cells were co-electroporated as described previously (36) with 5ug J6/JFH-1(p7-Rluc2a) RNA or related point mutant viral RNAs, 60 pmol of test or control siRNAs and 60 pmol of anti-miR-122. In all samples in an experiment the amount of small RNAs added per sample was equivalent, and if necessary siControl was added to balance the amount of small RNA (13). Cells were harvested 2 hours, 24 hours, 48 hours and 72 hours post-electroporations and assayed for Rluc expression. HCV replication was assessed based on Rluc expression and was normalized to a positive control sample in which replication of J6/JFH-1(p7-Rluc2a) RNA was supported by endogenous cellular miR-122 (Endo miR-122).

### Generation of Ago2/ DROSHA knockout Huh-7.5 cells

To generate the double knockout Huh 7.5 cells, we used CRISPR-Cas9 gene editing system to knockout Ago2 in DROSHA knockout Huh 7.5 cells provided to us by Dr. Charlie Rice (34). Three synthetic guide RNAs (AAUACCUGUUAACUCUCCUC-140585131; UAAUUUGAUUGUUCUCCCGG-140585231, GGCGCAGGAGGUGCAAGUGC-140585310) were designed using the Synthego knockout design tool (https://design.synthego.com/#/) in such a way that the guide RNAs would result in a frameshift deletion in the early region of the exon 2 in the Ago2 gene. DROSHA knockout cells were transfected with the TrueCut™ Cas9 Protein v2 (Invitrogen, Thermo Fisher Scientific, Vilnius, Lithuania) and the guide RNAs (Synthego CRISPRevolution EZkit, Redwood City, USA) using the Lipofectamine™ CRISPRMAX™ Cas9 Transfection Reagent (Invitrogen, Thermo Fisher Scientific, Carlsbad, USA) according to the manufacturer’s protocol (Quick Reference: Invitrogen, Thermo Fisher Scientific, Lipofectamine CRISPRMAX Transfection Reagent Pub. No.: MAN0014545). 48 hours post transfection, the cells were passaged, and a portion of the cells collected to test the knockout efficiency. Knockout efficiency was assessed by sanger sequencing of a PCR product that amplified the Crispr targeted region of the Ago2 gene, generated using primers-forward: 5’ATTCATGCTGCCTCATCTCTCC3’ and reverse: 5’CGGAAGAAGGTATGAGGCAA3’. PCR was performed using the PfuUltra II Fusion High-fidelity DNA polymerase (Agilent, California, USA) using genomic template DNA extracted using QuickExtract DNA extraction solution (Epicentre/ Lucigen, Wisconsin, USA). DNA was prepared by harvesting the cells in the QuickExtract DNA extraction solution and heating the samples at 65⁰C for 15 min, 68⁰C for 15 min and 98⁰C for 10 min. The ABI file obtained after sanger sequencing was examined for indels on the Synthego ICE tool (https://ice.synthego.com/<u>#/</u>) and the knockout efficiency was found to be 90%. An array dilution method was used to isolate single cell colonies and condition media was used to encourage the growth of the single clones. Condition medium was obtained by collecting the medium used for growing the knockout pool of cells, filtered using a 0.22µm filter and stored at −20⁰C prior to use. Successful knockout of both Ago2 alleles in the DROSHA knockout cells was identified based on analysis for the loss of siRNA knockdown phenotype based on HCV replication promotion by 5’ UTR annealing siRNAs and confirmed by western blot and genome sequencing.

### Phenotypic analysis of DROSHA/Ago2 knockout cells

To confirm knockout of Ago2 in the DROSHA/Ago2 KO cells we assessed the ability of the cells to support HCV replication promotion by si18-36, as compared to DROSHA KO wild type cells in which the siRNA will knock-down and thus fail to promote HCV (Figure 8A). Isolated clones of putative DROSHA/Ago2 double knockout cells were seeded in 24-well plate 24 hours prior to transfection with 1 ug of J6/ JFH-1(p7-Rluc2a) RNA and 12 pmol small RNAs. miR-122 was used as positive control and siControl was used as negative control. Transfections were performed using Lipofectamine 2000 (Invitrogen, Carlsbad, USA) as per manufacturer’s protocol. Cells were harvested 48 hours post transfection and Rluc expression was measured.

### Western blot

Knockout of Ago2 was confirmed by western blot analysis for the expression of Ago2 (Figure 8B), (Supplementary Figure 2C). Putative DROSHA/Ago2 knockout cells were treated with 1x SDS lysis buffer (with 1% 1M DTT) and heated at 95⁰C for 5min. The proteins were then separated using a 7.5% SDS-PAGE gel and transferred to a nitrocellulose membrane (GE healthcare Lifesciences, Amersham Protran 0.45 NC membranes, Freiburg, Germany). The membrane was blocked with 5% skimmed milk (BD Difco) and probed with 1:1000 diluted primary anti-Ago2 rat monoclonal antibody clone 11A9 (Millipore Sigma/ Merck KGaA MABE 253, Darmstadt, Germany), 1:25000 diluted primary mouse monoclonal anti-beta actin antibody (AC-15) (Abcam ab6276, Cambridge, USA) and subsequently with 1:40000 diluted secondary peroxidase conjugated AffiniPure Goat Anti-Rat IgG (H+L) (Jackson Immunoresearch 112-035-003, West Grove, USA) and 1: 25000 diluted secondary HRP conjugated Goat anti-mouse IgG (H+L) (BioRad, Mississauga, Canada). The blot was developed using Clarity Western ECL substrate (BioRad, Mississauga, Canada) and imaged with BioRad ChemiDoc MP Imaging system.

### HCV translation assay

To assess siRNA promotion of HCV translation, DROSHA/Ago2 KO Huh-7.5 cells were co-electroporated with 5ug of non-replicative mutant HCV genomic RNA, J6/JFH-1(p7-Rluc2a) GNN, 1ug of control T7 Fluc mRNA, and 60 pmol of test siRNA. As positive control J6/JFH-1(p7-Rluc2a) GNN was electroporated with miR-122, since the DROSHA/Ago2 KO Huh-7.5 cells lack miR-122 expression. As a negative control, J6/JFH-1(p7-Rluc2a) GNN was electroporated with siControl. Cells were harvested 4 hours post-electroporations and assayed for Rluc and Fluc expression.

### Luciferase assay

Luciferase expression was measured by using Firefly, *Renilla*, or Dual luciferase kits (Promega, Madison, USA) as suggested by the manufacturer’s protocols. Cells were washed once in Dulbecco’s phosphate-buffered saline then lysed with 100 ul of passive lysis buffer. 10 ul of the cell extract was mixed with the appropriate luciferase assay substrate and light emission was measured by using a Glomax 20/20 Luminometer (Promega, Madison, USA).

### RNA purification

Cells were harvested into 1 ml of Trizol and total cellular RNA was isolated using the manufacturer’s provided protocol (Life Technologies, Burlington, Canada).

### HCV genome stabilization assay and northern blots

To assess the impact of miR-122 and the siRNAs on HCV genome stability we assessed the amount of non-replicative HCV RNA present in cells at various times post-electroporation using northern blot analysis (Figure 9), (Supplementary Figure 2A and B). For each assay, 32 × 10^6^ DROSHA/Ago2 knockout Huh-7.5 cells were electroporated (in 4 cuvettes) with 40 μg of HCV J6/JFH-1(p7-Rluc2a) GNN RNA and 240 pmol of one of the small RNAs (si15-33, si19-37, si26-44, or si27-45) or miR-122 (as a positive control) or siControl (as a negative control). Cells from the 4 electroporation cuvettes were pooled and plated onto four 10 cm plates. Cells for the 0 min time point were harvested immediately after electroporation, and the others were incubated at 37°C and harvested at 30 mins, 60 mins and 120 mins post-electroporation. Total cellular RNA was harvested using Trizol as recommended by the manufacturer (Life Technologies, Burlington, Canada), and 10 ug were separated on an 0.8% agarose gel and transferred to a Nylon membrane (GE Healthcare Limited, Buckinghamshire, England) as described previously (36). The transferred RNAs were crosslinked to the membrane using a UV crosslinker (Spectrolinker XL-1000) at X100 μJ/cm^2^ for 12 seconds and cut in half to probe for HCV RNA and GAPDH separately. The radioactive DNA probes used were prepared using Prime-a-Gene Labeling System kit (U1100, Promega, Madison, WI, USA), and radiolabeled dCTP (PerkinElmer, Boston, USA). The probes were generated from a 3 kbp BamHI-to-EcoRV DNA fragment of the pJ6/JFH-1 RLuc (p7-RLuc2A) plasmid or a 1.3 kb cDNA fragment of human GAPDH. Radioactive bands were detected by exposing the membranes overnight on a phosphorscreen and scanned using a Phosphoimager (Typhoon, GE Healthcare Life Sciences, Mississauga, Canada). Band signal intensities were quantified using Image Studio Lite version 5.2.5.

### RNA structure prediction analysis

RNA structure predictions were done using the RNA prediction software ‘RNA structure’ available from the website by the Matthews lab at https://rna.urmc.rochester.edu/index.html. (37). Single RNA structure predictions were performed using algorithm ‘fold’ and structure predictions of two interacting RNA molecule were predicted using algorithm ‘bifold’. Dot-bracket files for the five lowest free energy structures were generated using the RNA fold command in ‘RNAstructure’ and RNA images were generated from them using VARNA (VARNA GUI applet) (38).

### Statistical analysis

All data are displayed as the mean of three or more independent experiments, and error bars indicate standard deviation of the mean. Where appropriate, one-way ANOVA was performed using Graph Pad Prism version 8.3 for MacOS (San Diego, USA, www.graphpad.com). In graphs, statistical significance is indicated as follows: *P<0.0332; **P< 0.0021; ***P < 0.0002; ****P < 0.0001.

## RESULTS

### siRNA annealing to nucleotides between 15 to 44 on 5’UTR promotes virus replication

Annealing of miR-122 to two complementary sequences on 5’UTR of the HCV genome is required for detectible HCV replication in cell culture (27, 39). In our previous work we showed that HCV replication was promoted efficiently by 5’ UTR targeting siRNAs when siRNA directed cleavage activity was abolished by using Ago2 knockout cells (Ago2 KO) (13). Thus, replication promotion does not require the specific annealing pattern generated by miR-122 or binding of two small RNA copies, but we hypothesized that it may be impacted by the annealing location. To test this hypothesis, we determined the range of genome locations to which small RNA annealing can promote HCV replication. First, we designed and tested an array of siRNAs with target sequences that walk the 5’UTR between nucleotides 10 to 47 at single nucleotide resolution (Figure 2A). Each siRNA was 19 nucleotides long and contained two 3’ UU (uracil) overhangs to ensure RISC loading (Supplementary Table 3). The siRNAs were named based on the 19 nucleotide positions on the HCV genome to which they bind, from si10-28 to si29-47 (Figure 2A). To confirm RISC loading, the siRNAs were tested for their ability to knockdown gene expression in a suppression assay (Figure 2B). For suppression assays, we transfected cells with a plasmid, pFluc JFH-1 5’UTR, that expresses an mRNA encoding Fluc having the HCV miR-122 binding region/siRNA target sites in its 3’ UTR. This plasmid was co-transfected in miR-122 KO Huh 7.5 cells (cells that express Ago2), with a test siRNA and knockdown was measured based on Fluc expression compared to cells transfected with a control siRNA (siControl). Fluc expression levels were assessed relative to Rluc expression from a transfection control plasmid, pRL-TK. We observed that siRNAs binding from nucleotides 14 onwards were significantly (p value <0.0001) able to suppress Fluc expression at different levels compared to siControl, implying that they were actively incorporated into the RISC complex (Figure 2C) (Supplementary Table 1). Most siRNAs that bound within SLI did not knockdown Fluc and thus did not enter RISC, likely due to the hairpin structure formation. Inactive siRNAs were omitted from further analyses. After confirming RISC incorporation, we determined the ability of the siRNAs to promote HCV replication in replication assays. For these assays we electroporated Ago2 KO Huh 7.5 cells with J6/JFH-1(p7-Rluc2a) RNA, anti-miR-122, an antagonist of endogenous miR-122, and a test siRNA, and HCV replication was assessed 2 hours, 24 hours, 48 hours and 72 hours post-electroporation based on luciferase expression as a proxy for HCV replication (Figure 2D). We electroporated J6/JFH-1(p7-Rluc2a) RNA without anti-miR-122 as a positive control to measure HCV RNA replication induced by endogenous miR-122 (Endo miR-122). RLuc expression in this sample at 72 hours post-electroporation was deemed 100% and used to calculate relative luciferase levels in the rest of the samples. The negative controls (siControl) (Figure 2D) containing viral RNA, anti-miR-122 and siControl (an HCV non-targeting siRNA) confirmed abolition of HCV replication by the miR-122 antagonist. In other samples the addition of indicated siRNA reinstated HCV replication to the relative levels shown (Figure 2D), (Supplementary Table 1). An siRNA that anneals within the NS5B coding sequence (si6467) did not promote HCV replication but siRNAs binding between nucleotides 15 and 44 did. HCV replication was promoted most efficiently by si19-37 and was similar to replication induced by endogenous miR-122 (Endo miR-122), and less efficient replication induction was observed using siRNAs that annealed to locations moving away from nucleotides 19-37 in either direction (Figure 2D and F). The 5’ boundary to which siRNAs can promote HCV was nucleotide 15. Si14-32 did not promote detectable HCV replication and siRNAs that anneal nearer to the 5’ UTR did not enter RISC and thus could not be tested (Figure 2C). The 3’ boundary of the region to which siRNA annealing promoted HCV RNA was nucleotide 44 and si26-44 was the last siRNA to promote virus replication (p value <0.0001). si26-44 promoted HCV replication; however, si27-45 and those that bound further in the 3’ direction did not. Thus, either annealing to nucleotide 26 was essential for promotion, or siRNA annealing to nucleotide 45 and onwards was detrimental. To distinguish between these possibilities, we designed two additional siRNAs, si26-45 and si27-44. Both siRNAs were active in suppression assays, however only si27-44 promoted HCV replication (Figure 2E), (Supplementary Table 1). This indicated that annealing to nucleotide 45, inhibited promotion activity and annealing to nucleotide 26 is not essential (Figure 2E). Thus, we have defined the 3’ boundary of the region to which small RNA annealing can promote HCV replication as nucleotide 44.

### siRNA annealing to the HCV IRES do not promote or inhibit HCV replication

To determine whether siRNAs binding on other regions of HCV genome can promote virus replication, we designed siRNAs using online software, i-score, to determine best target sites within the HCV IRES region (31). We designed 6 siRNAs that bind to various sites on the IRES regions namely si38-56, si42-60, si73-91, si88-106, si317-338 and si339-357 (Figure 3A). We validated the knockdown activity of these siRNAs to confirm their incorporation into RISC, and, 5 out of 6 showed suppression activity (Supplementary Figure 1A). However, none of the active siRNAs promoted replication in HCV replication assays (Figure 3B).

**Figure 3:**
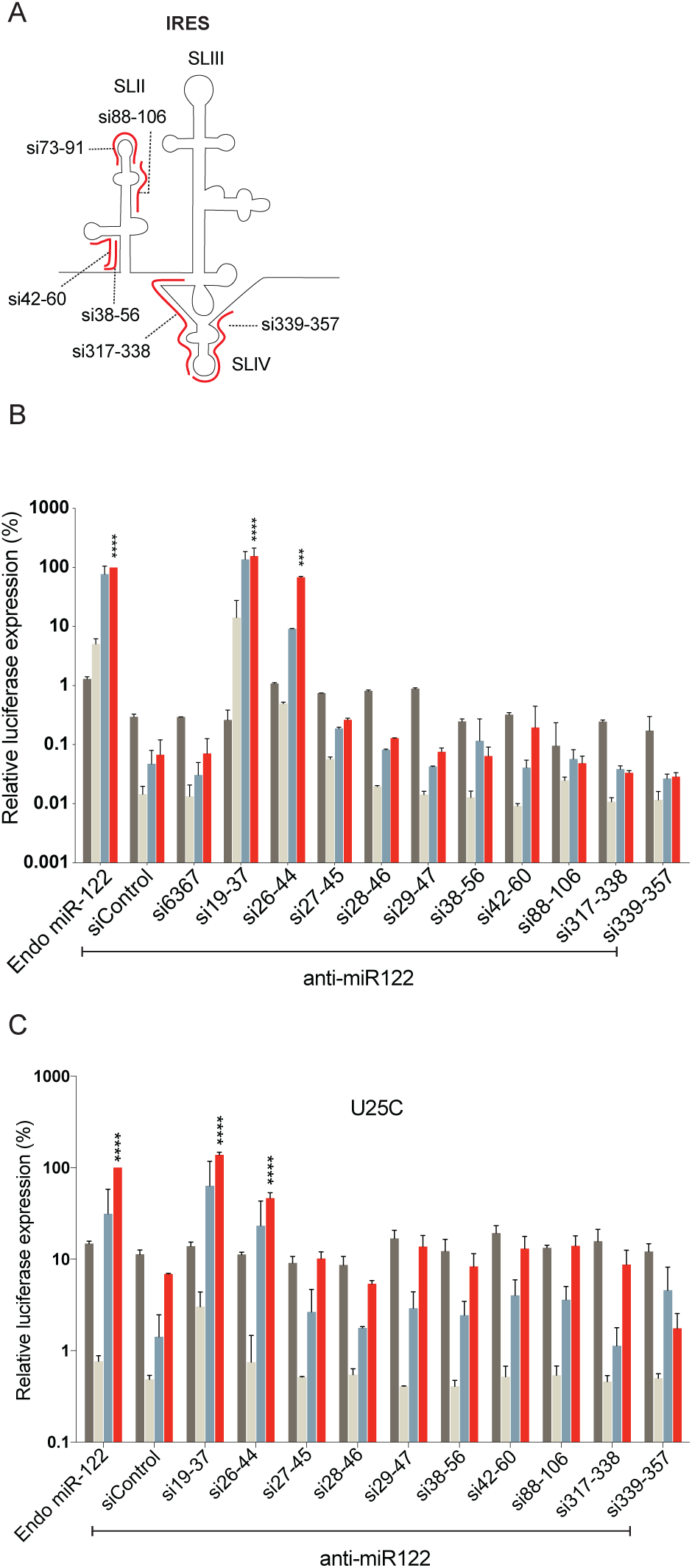
The pro-and antiviral activity of siRNAs binding to the HCV IRES. (A) A diagram of the HCV IRES including stem loops II, III and IV and the locations of annealing of six siRNAs that target the IRES are shown in red. (B) The pro-viral activity of the siRNAs was assessed using replication assays in which the activity of miR-122 is antagonized by anti-miR-122. Ago2 knockout cells were electroporated with HCV J6/JFH-1(p7-Rluc2a) RNA, anti-miR-122 and the indicated siRNA and harvested at 2 hours (grey bars), 24 hours (light grey bars), 48 hours (blue bars) and 72 hours (red bars) post-electroporation. (C) The proor anti-viral effects of the indicated siRNAs was assessed based on their influence on miR-122-independent replication of a U25C mutant J6/JFH-1(p7-Rluc2a) similar to the experiments described in B. For B and C, HCV replication was measured based on Rluc expression and is presented as % relative to Rluc expression from HCV RNA supported by endogenous miR-122 at 72 hours post electroporation (Endo miR-122). The data are the average of at least 3 independent experiments and error bars represent the standard deviation. Statistical significance was determine using one-way ANOVA on the relative 72 hour values where, *P<0.0332; **P< 0.0021; ***P < 0.0002; ****P < 0.0001.

Small RNAs that anneal within the IRES and downstream of nucleotides 45, and thus within SLII of the IRES did not promote detectible HCV replication. We hypothesized that these siRNAs may have failed to promote HCV because they anneal to, and interfere with the activity of the IRES and thus inhibit HCV replication. To test this hypothesis, we assessed inhibition of HCV by the IRES targeting siRNAs. For these assays we used a mutant HCV RNA, U25C, that can replicate independent from miR-122, and thus replicated in Ago2 knockout cells even when miR-122 is antagonized. This assay was used instead of miR-122-promoted replication to eliminate the possible influence of annealing competition between miR-122 and the siRNAs tested. Using this assay, we observed no HCV replication promotion or inhibition by the IRES-binding siRNAs, while replication was promoted by the positive control siRNA, si19-37 (Figure 3C). Thus, our data suggests that IRES-binding siRNAs do not promote or inhibit HCV replication.

### siRNA annealing to the alternative predicted miR-122 binding sites do not promote HCV replication

Other potential miR-122 binding sites have been reported in the NS5B coding region and the 3’UTR region of HCV genome (40–43). We therefore also assessed whether siRNAs binding on any of these potential miR-122 binding sites affected HCV replication (Figure 4). We started by designing siRNAs binding to 7 predicted miR-122 binding sites in NS5B and 3’UTR region. For convenience we named these siRNAs according to the region they bind and in serial order, namely siNS5B1, siNS5B2, siNS5B3, siNS5B4, si3’UTR1, si3’UTR2 and si3’UTR3 (Figure 4A and B). In order to test these in our suppression assay, we replaced the 5’UTR with NS5B-3’UTR regions in suppression plasmid to have target sites for our siRNAs. We observed that each of seven siRNAs were able to suppress Fluc expression suggesting they all entered RISC (Supplementary Figure 1B). To assess their role in HCV replication, we performed HCV replication assays and observed that Rluc expression after addition of the NS5B and 3’ UTR targeting siRNAs was comparable to the negative controls (siControl and si6367) (Figure 4C). Thus, siRNA binding to other predicted miR-122 binding sites did not promote virus replication. Overall our data show that small RNAs that bind between nucleotides 15-44 on the HCV 5’ UTR promote HCV replication.

**Figure 4:**
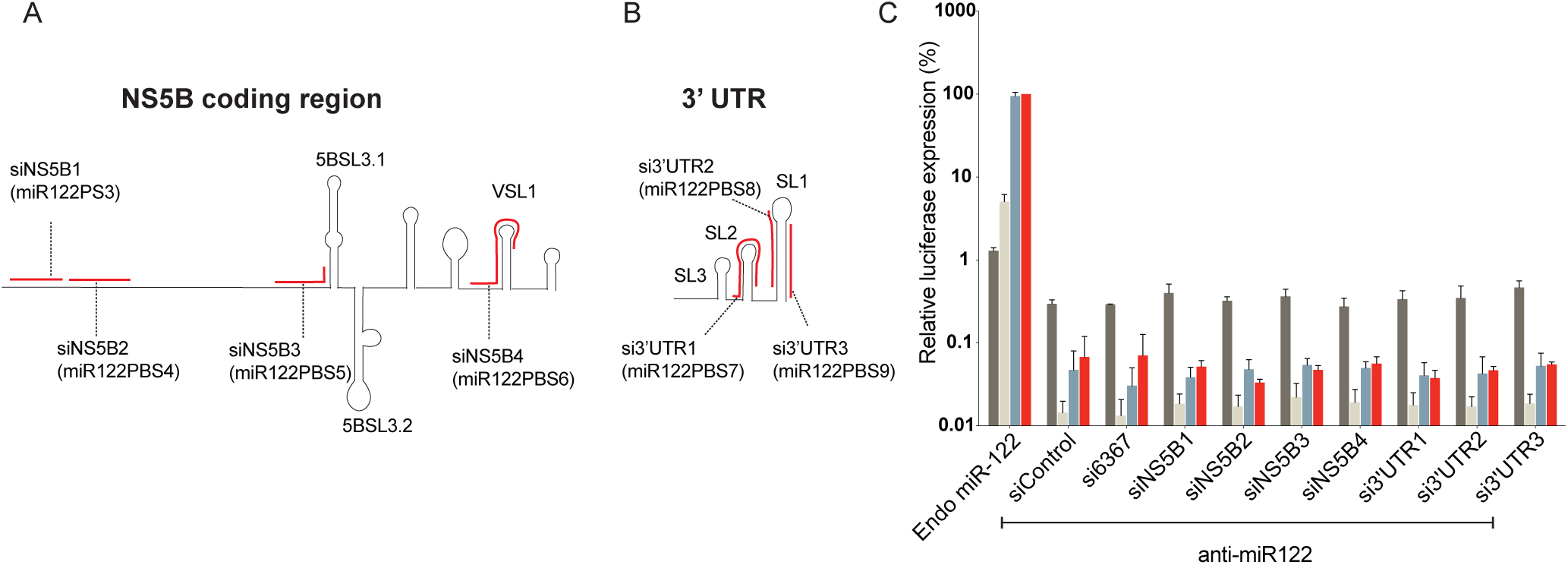
The influence on HCV replication of siRNAs that binding at other predicted miR-122 binding. (A, B) Diagrammatic representation of HCV NS5B coding region and 3’UTR region secondary structures. The figure shows siRNAs (red) binding to other predicted miR-122 binding sites in NS5B coding region (left) and 3’UTR region (right). (C) The activity of the siRNAs was assessed using replication assays in which the activity of miR-122 is antagonized by anti-miR-122. Ago2 knockout cells were electroporated with HCV J6/JFH-1(p7-Rluc2a) RNA, anti-miR-122 and the indicated siRNA and harvested at 2 hours (grey bars), 24 hours (light grey bars), 48 hours (blue bars) and 72 hours (red bars) post-electroporation. HCV replication was measured based on Rluc expression and is presented as % relative to Rluc expression from HCV RNA supported by endogenous miR-122 at 72 hours post electroporation (Endo miR-122). Data represent the average of 3 independent experiments and error bars represent the standard deviation. Statistical significance was determine using one-way ANOVA on 72-hour values where, *P<0.0332; **P< 0.0021; ***P < 0.0002; ****P < 0.0001.

### Small RNAs that promote virus replication are predicted to induce the canonical 5’ UTR RNA structure

We and others hypothesized that the pro-viral activity of miR-122 was mediated by annealing induced RNA structural changes to the HCV 5’ UTR that promote the formation of the canonical 5’ UTR IRES (Figure 1 A and B) (12, 13, 24). In the presence of miR-122 the HCV 5’ terminal 117 nucleotides were predicted to form SLIIa and SLIIb, essential IRES elements, and SLI, an essential replication element (Figure 1A and C). However, in the absence of miR-122, the 5’ UTR RNA is predicted to form SLI and an alternative structure instead of SLII and thereby would fail to form an active HCV IRES (Figure 1B). Since siRNAs annealing to different locations promoted virus replication with different efficiencies, we speculated that replication promotion may correspond with the ability of an siRNA to induce the canonical HCV 5’ UTR RNA structure. To test this, we used an online RNA secondary structure prediction tool, ‘RNAstructure’ to predict the structure of the 5’ 117 nucleotides of the HCV genome induced by annealing of siRNAs that promote efficient, intermediate, or no HCV replication. The first six lowest delta free energy predicted structures induced by annealing of each siRNAs are shown in Figure 5. We observed that siRNAs that promote efficient virus replication, si17-35, si19-37, si22-42 and si24-42 were predicted to induce the canonical HCV structure one or many times of the six predicted structures (Figure 5), (Supplementary Tables 1 and 2). We also found that the canonical structure was induced by annealing of si26-44 and si27-44 which mark the boundary for replication promotion. However, the canonical structure was not observed with siRNAs si26-45 and si27-45, and whose annealing did not promote HCV replication. This was also observed for si14-32, which did not promote virus replication and was predicted to form an additional non-canonical loop near SLIIa (Figure 5, si14-32, Structure no. 4), and si28-46 which induced a structure in which a stem of SLIIa was absent (Figure 5), (Supplementary Table 2). Si15-33 showed intermediate replication promotion and also failed to induce SLI (Figure 5), (Supplementary Table 2). Thus, this data suggests that formation of the complete canonical 5’ terminal RNA structure, including SLI, SLIIa and SLIIb by small RNA annealing is required for optimal HCV replication promotion, and that an incomplete canonical structure impairs viral replication.

**Figure 5:**
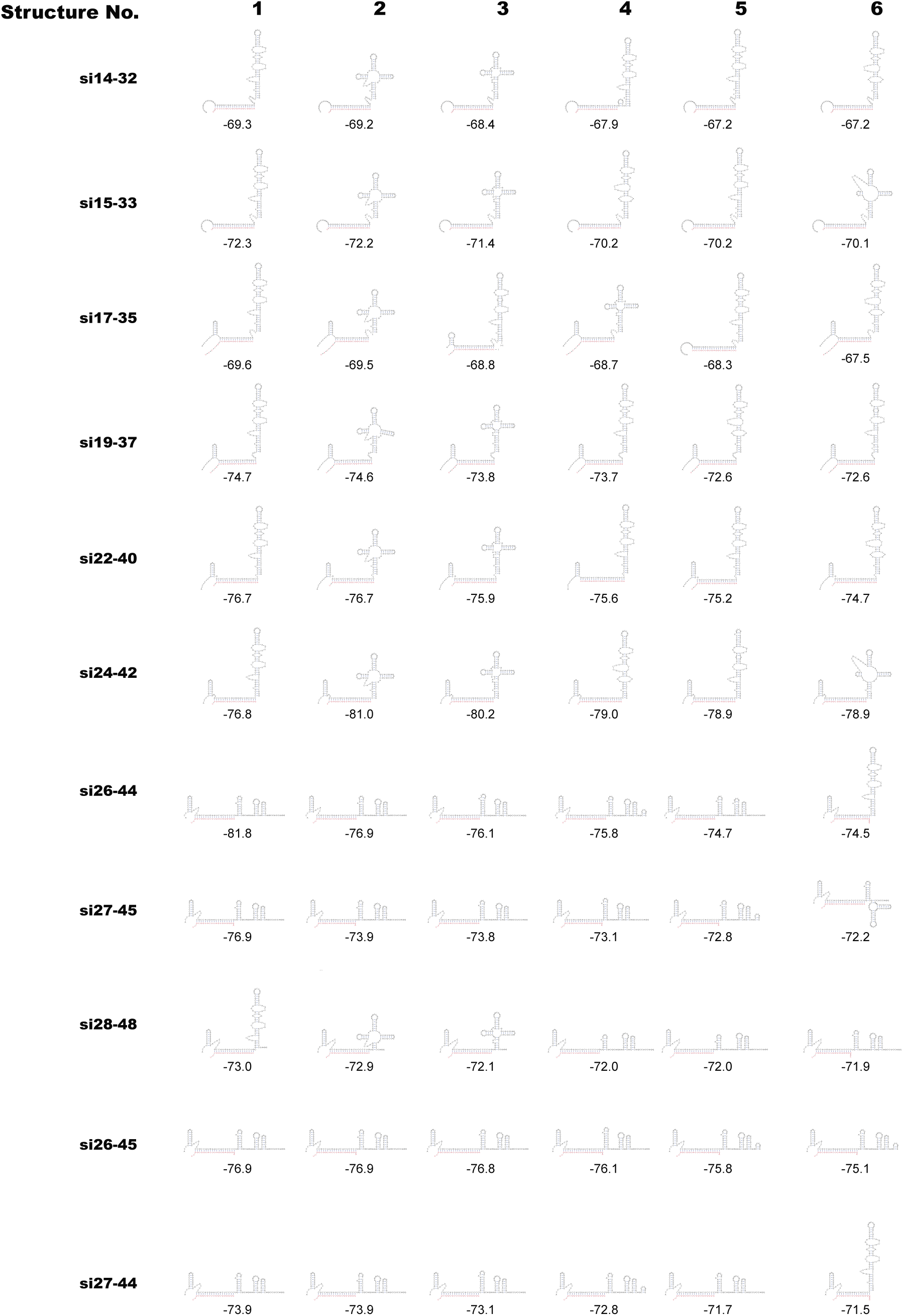
5’ UTR RNA structure predictions following annealing of the siRNAs that promote and do not promote HCV replication. Structure predictions of the HCV 5’ terminal 117 nucleotides with annealing of the indicated siRNA was performed using online software RNAstructure, specific algorithm bifold. The first six lowest delta free energy structures obtained from predictions with each siRNAs are shown. The delta free energy calculated by the software is shown below each structure.

### Binding of siRNAs to 15 nucleotides is a minimum requisite to promote efficient small RNA dependent HCV replication

Next we wanted to define the minimum annealing requirements for the pro-viral activity of small RNA annealing (Figure 6). To determine this, we designed siRNA analogues of the most efficient siRNA, si19-37, but having sequence matches ranging from 7 to 17 nucleotides on the 3’ or 5’ ends (Figure 6A). For example: si19(21–37) is 19 nucleotides long with 17 nucleotides (21-matching the HCV 5’ UTR (Figure 6A). Similarly, si19(23–37), si19(25–37), si19(27–37), si19(27–37), si19(29–37) and si19(31–37) have between 15 and 7 matching nucleotides. si19(21–37), si19(23–37) promoted HCV replication as efficiently as si19-37 and the others did not promote at all (Figure 6B). From the 3’ end we generated siRNAs with sequence matches of 16, si19(19–35), 14, si19(19–33), and 12, si19(19–31) matches (Figure 6A) and only si19(19–35) promoted HCV replication (Figure 6C). Suppression assay of all si19-37 analogs showed that siRNA knockdown activity decreased with fewer annealing nucleotides; however, those that promoted replication were all active (Supplementary Figure 1C and D). Thus, with this we can conclude that an siRNA must anneal to at least 15 nucleotides of the HCV 5’ UTR to promote efficient replication. RNA structures induced by annealing of these siRNAs resembled the canonical structure, but delta free energy data suggest that the structures may be less stable (Figure 6D). Therefore, our data suggests that location specific siRNA must anneal with a minimum of 15 nucleotides to induce the canonical structure and promote efficient virus replication.

**Figure 6:**
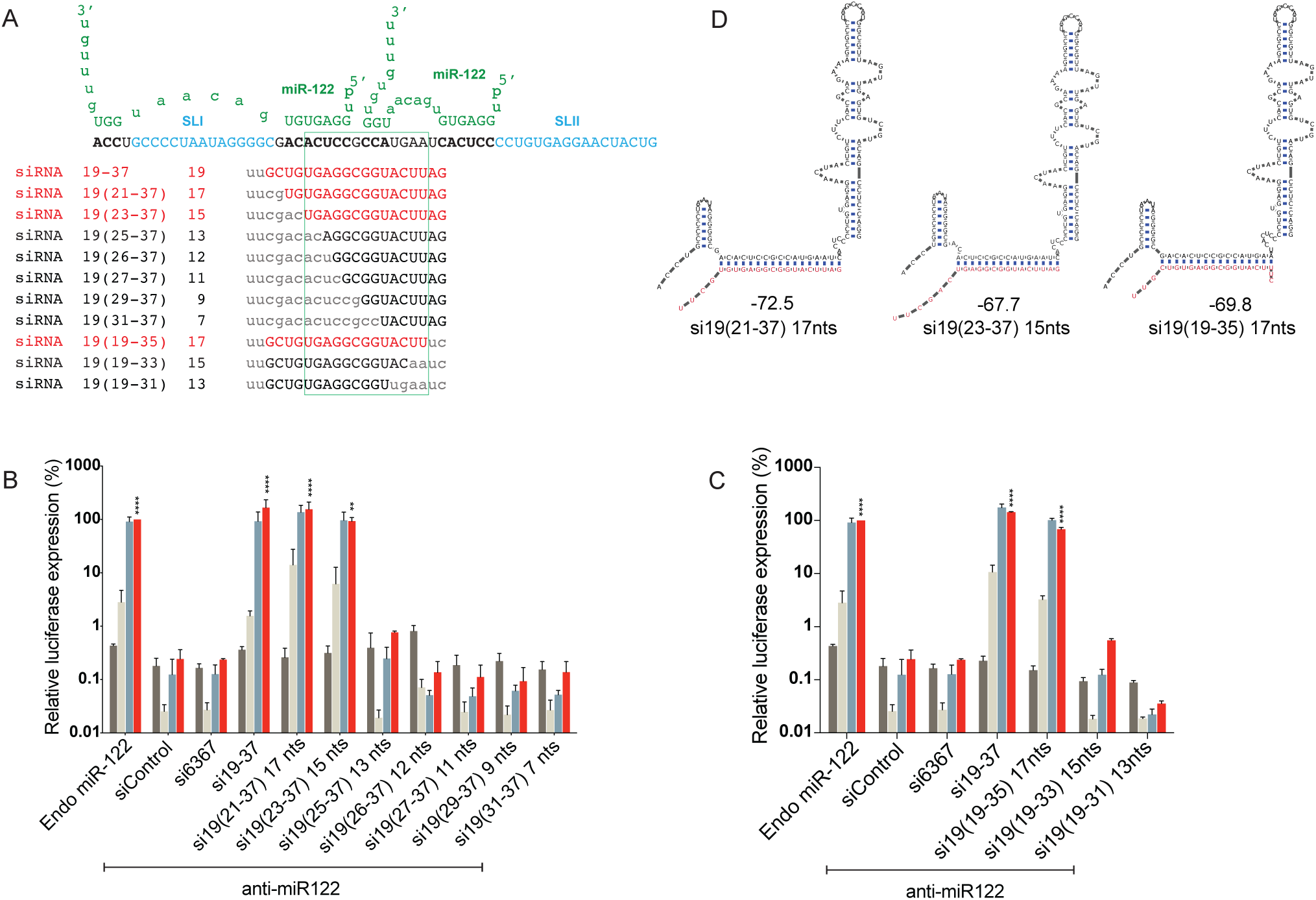
The minimum annealing requirements of siRNAs with si19-37 analogues. (A) Diagrammatic representation of first 55 nucleotides of HCV 5’ UTR are shown interacting with 2 copies of miR-122 (green). Sequences of si19-37 analogues at their binding positions are shown and the number of annealing nucleotides for each analogue siRNA is indiacated. siRNAs in red promote HCV replication and siRNAs in black do not promote HCV replication. Small lettered characters in siRNAs show mismatched nucleotides that do not bind on HCV RNA (grey) while capital lettered characters are nucleotides that bind on the RNA. The green box shown around nucleotides 23-35 which were common in case of all siRNAs that promote HCV replication. (B and C) The activity of the si19-37 analogues was assessed using replication assays in which the activity of miR-122 is antagonized by anti-miR-122. Ago2 knockout cells were electroporated with HCV J6/JFH-1(p7-Rluc2a) RNA, anti-miR-122 and the indicated siRNA and harvested at 2 hours (grey bars), 24 hours (light grey bars), 48 hours (blue bars) and 72 hours (red bars) post-electroporation. HCV replication was measured based on Rluc expression and is presented as % relative to Rluc expression from HCV RNA supported by endogenous miR-122 at 72 hours post electroporation (Endo miR-122). Data represent

### siRNAs binding to 5’ terminus (nts 1-3) enhances HCV replication, but the generation of a small RNA overhang does not

Previous reports showed that replication promotion by miR-122 was enhanced by annealing of miR-122 to nucleotides on the 5’ terminus of HCV and by the generation of a 3’ overhang (20). To assess the impact of siRNA binding to the 5’ terminus (nucleotides 1-3) we compared HCV replication promotion by a small RNA that binds to the 5’ terminus with one that does not (Figure 7). We designed an siRNA, si1-3--21-36, that binds to 21-36 and to the 5’ terminal 3 nucleotides and found that it promoted replication about 4 fold more efficiently than si1-3mm--21-36, that also binds to nts 21-36 but not the 5’ terminus (Figure 7A and B). Both siRNAs were active in our suppression assays (Supplementary Figure 1E). Indeed, HCV replication promoted by si1-3--21-36 was around 2-fold higher than the most efficient siRNA, si19-37, identified in Figure 2D. Further, the canonical 5’ UTR RNA structure was induced regardless of 5’ end annealing. This supports that end annealing is not required for pro-viral siRNA annealing but suggests that it has a positive effect on virus replication (Figure 7C).

**Figure 7:**
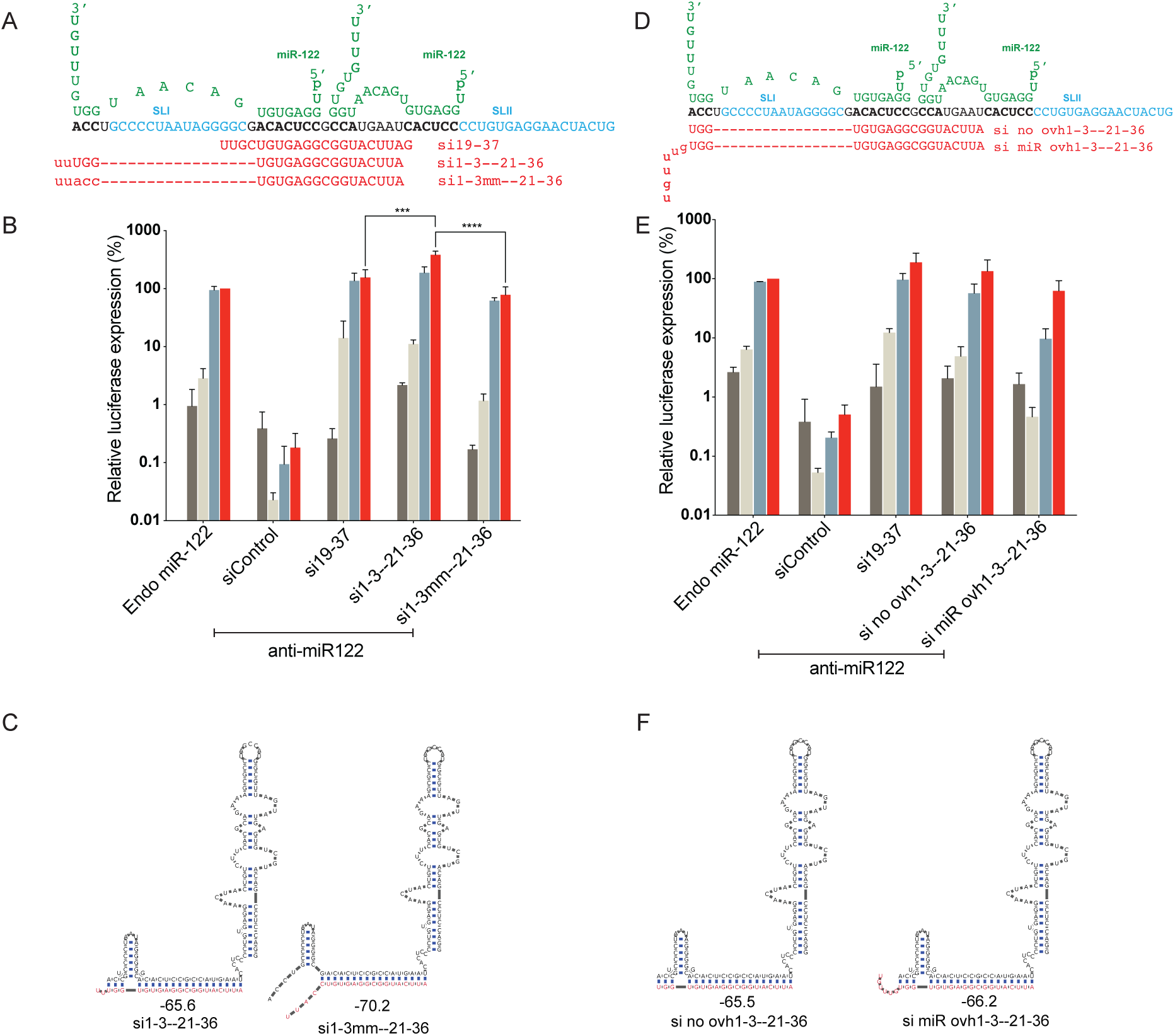
HCV replication promotion with siRNAs binding on HCV 5’ terminus and an siRNA generated 3’ overhang. (A and D) Diagram of the first 55 nucleotides of HCV 5’ UTR interacting with 2 copies of miR-122 (green). miR-122 binding sites on HCV genome are shown in bold characters. Uppercase letters represent nucleotides in the siRNAs that anneal to the HCV RNA and lowercase letters represent nucleotides that do not. (A). Sequences of si19-37, si1-3--21-36 and si1-3mm--21-36 are shown. (D). Schematic representation of si no ovh1--21-36 and si miR ovh1-3mm-21-36 binding on HCV genome. (B and E). Graphs showing HCV replication promotion by siRNAs that bind or do not bind to the 5’ terminus and with siRNAs that do or do not generate a 3’ overhang. Ago2 knockout cells were electroporated with HCV J6/JFH-1(p7-Rluc2a) RNA, anti-miR-122 and the indicated siRNA and harvested at 2 hours (grey bars), 24 hours (light grey bars), 48 hours (blue bars) and 72 hours (red bars) post-electroporation. HCV replication was measured based on Rluc expression and is presented as % relative to Rluc expression from HCV RNA supported by endogenous miR-122 at 72 hours post electroporation (Endo miR-122). Data represents the average 3 independent experiments and error bars represent the standard deviation. Statistical significance was determine using one-way ANOVA on 72-hour values where, *P<0.0332; **P< 0.0021; ***P < 0.0002; ****P < 0.0001. (C and F) RNA structure predictions of the HCV 5’ terminal 117 nucleotides with annealing of siRNAs that bind or do not bind to 5’ terminus and with siRNAs that do or do not generate a 3’ overhang. Predictions are based on bifold in the online software RNAstructure and the delta free energy calculated by the software are shown below each structure.

The annealing of miR-122 to binding site 1 on the HCV genome generates a 7-nucleotide overhang of the HCV 5’ terminus that was reported to contribute to the efficiency of replication of HCV with miR-122 (20). Since the siRNAs used in this study only consisted of 2 UU overhangs, we knew that a long miR-122 like overhang was not essential for replication promotion, but we wanted to test if having such an overhang contributes to replication. To test this, we assessed replication promotion by an siRNA that has a miR-122-like overhang, ‘si mir ovh1-3--21-36’, with one that does not, ‘si no ovh1-3--21-36’ (Figure 7D). Both siRNAs were active in our suppression assays (Supplementary Figure 1E). Further, both siRNAs are predicted to induce formation of canonical predicted HCV structures (Figure 7F) but we found that the miR-122-like overhang did not enhance replication promotion, and in fact, decreased HCV replication efficiency (Figure 7E). This experiment confirmed that generation of a 5’ overhang is not required for and may hinder HCV replication.

### siRNA promotion of virus replication correlates with promotion of virus translation

Two confirmed functions of miR-122 are promotion of HCV translation and stabilization of the viral genome, however, the relative contributions of each to HCV life-cycle promotion are unknown (27, 39, 44). We hypothesized that if stimulation of translation or genome stabilization is a key mechanism by which miR-122 promotes virus replication then the ability of an siRNA to promote replication will correlate with its ability to stimulate translation or stabilize the viral genome. To test this hypothesis, we assessed siRNA stimulation of translation and genome stabilization by panels of siRNAs that promote HCV with varying efficiencies.

To assess siRNA translation promotion, we measured the ability of miR-122 and an array of siRNAs to promote translation of a non-replicative HCV J6/JFH-1(Rluc2a) GNN RNA in DROSHA/Ago2 double KO cells. DROSHA/Ago2 double KO cells were generated to provide a background that lacked both miR-122 expression and Ago2 associated siRNA cleavage activity and allowed us to remove the miR-122 antagonist from our assays. DROSHA/Ago2 double KO cells were generated from DROSHA knockout cells using Crispr/Cas9 and the knockout of Ago2 was confirmed based on the ability of the cells to use si18-36 to promote instead of knockdown HCV replication (Figure 8A) and by western blot analysis showing abolished Ago2 expression. (Figure 8B), (Supplementary Figure 2C). To assess HCV translation promotion by the siRNAs, cells were electroporated with viral RNA and siRNAs and translation efficiency was measured based on Rluc expression vs a co-electroporated Fluc mRNA control (Figure 8C). By using an array of siRNAs that promote replication with different efficiencies we found that the levels of translation stimulation correlated with their ability to promote replication (Figure 8C and D), (Supplementary Table 1). siRNAs that promoted efficient HCV replication, si17-35, si18-36, si19-37, si22-40, and si24-42 also efficiently stimulated translation, and siRNAs that promoted HCV replication to a moderate level also promoted translation less efficiently. Finally, siRNAs that promoted HCV replication poorly displayed little or no translation stimulation (Figure 8C and D) (Supplementary Table 1). Our data thus suggests that translation stimulation is linked with small RNA promotion of HCV translation and supports the hypothesis that a primary mechanism of miR-122 promotion of HCV is by stimulating translation by inducing the formation of the HCV IRES.

**Figure 8:**
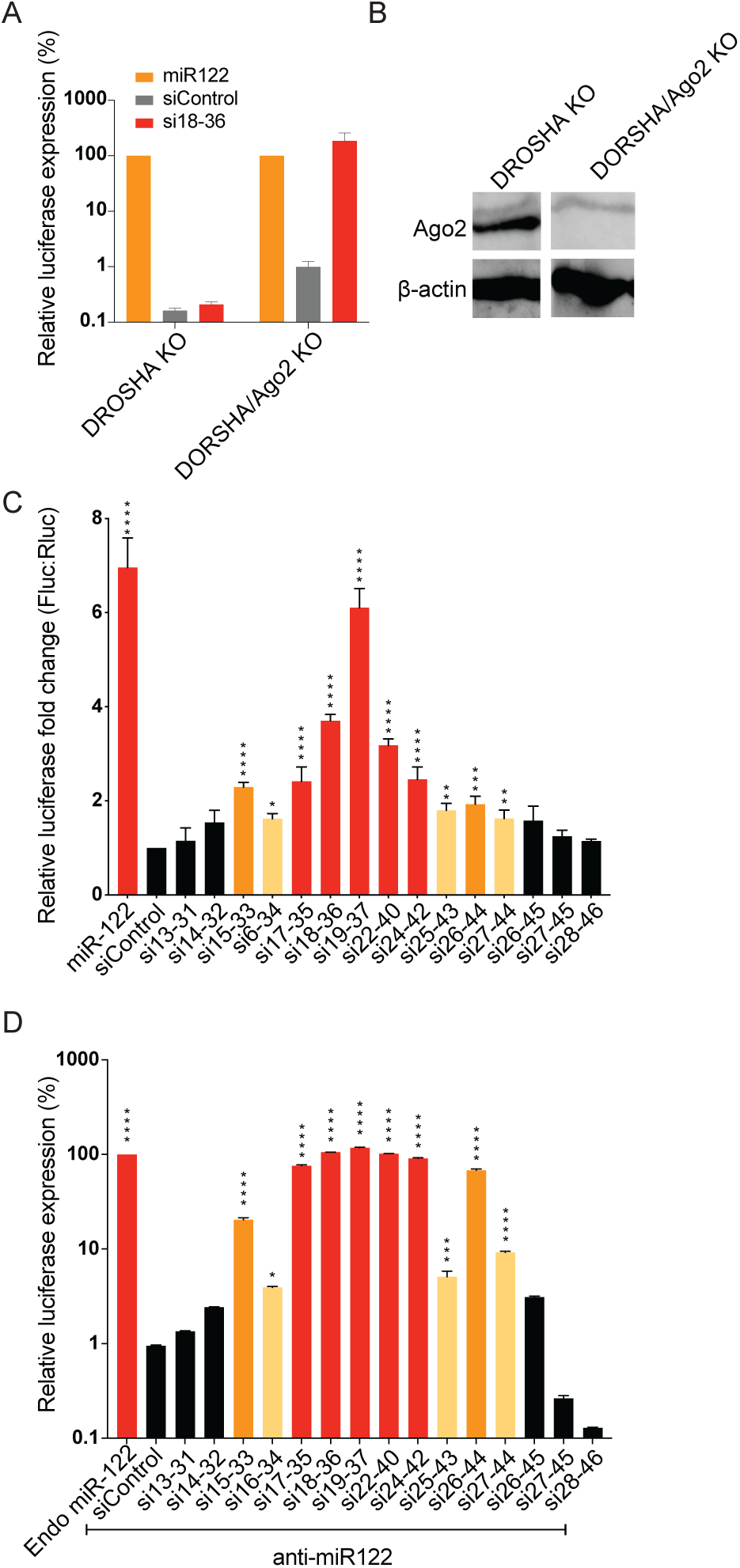
small RNAs annealing promotes translation of HCV. (A) Phenotypic analysis of DROSHA/Ago2 double KO cells. Graph showing HCV replication induction by siRNAs in DROSHA KO cells vs DROSHA/Ago2 double KO cells. (B) Western blot images showing the absence Ago2 protein in DROSHA/Ago2 double KO cells vs the presence of Ago2 protein in DROSHA KO cells. Uncropped blot is shown in Supplement Figure 2C (C) Transient translation assays performed in DROSHA/Ago2 double KO cells using non-replicative viral RNA, J6/ JFH-1(p7-Rluc2a) GNN, and the indicated siRNAs and an mRNA expressing Fluc as an electroporation control. Samples were harvested at 4 hours and translation was accessed based on Rluc expression vs the co-electroporated Fluc mRNA control. Data represent the average of at least 5 independent experiments and error bars represent the standard deviation. Statistically significant differences between siRNAs and siControl was assessed by one-way ANOVA, *P<0.0332; **P< 0.0021; ***P < 0.0002; ****P < 0.0001. (D) Graph showing replication promotion at 72 hours post infection by siRNAs used in the translation assays performed in Ago2 KO cells. (C and D) siRNAs that promoted HCV translation and replication efficiently are coloured red, moderately efficient ones are coloured orange and low efficient are coloured yellow. siRNAs that do not promote replication are coloured black.

### siRNA annealing-induced HCV genome stabilization is not sufficient to promote virus replication

In addition to stimulating HCV translation, miR-122 annealing also stabilizes the HCV genomic RNA by protecting it from cellular pyrophosphatases, Dom3Z and Dusp11, and the exonuclease Xrn1 (23). To test for a linkage between siRNA induced replication and virus genome stabilization we investigated HCV RNA genome stability in presence of miR-122, and four different siRNAs (Figure 9) (Supplementary Figure 2A and B). We chose si19-37 because it promotes efficient replication, two siRNAs, si15-33 and si26-44 that promote intermediate levels of replication, and one siRNA, si27-45 that does not promote replication. miR-122 was used as a positive control and siControl was used as a negative control. For the stability assays HCV J6/JFH-1(p7-Rluc2a) GNN RNA, a non-replicative HCV RNA and an siRNA (or miR-122, or control) were electroporated in DROSHA/Ago2 double KO cells and total RNA was harvested at 0min, 30 mins, 60 mins and 120 mins post-electroporation. To determine the half-life of HCV GNN RNA northern blots were performed (Figure 9). As expected, the half-life of the viral RNAs was extended by miR-122 annealing (Figure 9A), however, contrary to our expectations all siRNA stabilized the HCV genome, regardless of whether they promoted HCV replication or not (Figure 9, si27-45) (Supplementary Table 1). Further, no correlation between half-lives and the levels of RNA replication was observed (Figure 9A and B). This suggests that although small RNAs may stabilize the HCV genomic RNA this is not sufficient to promote HCV genome replication. Thus, stimulation of translation appears to be the essential role of miR-122 annealing in promotion of HCV replication, and genome stabilization, while stimulatory, is not sufficient alone.

## DISCUSSION

miR-122 binding to two sites on HCV 5’UTR is required for efficient HCV replication (Figure 2A). We previously reported a hypothesis that the pro-viral activity of miR-122 was mediated by annealing induced RNA structural changes to the HCV 5’ UTR to induce the formation of the canonical 5’ UTR IRES structure (13). We also showed that HCV replication was stimulated by siRNAs as efficiently as by miR-122 if their siRNA-directed cleavage activity was abolished by using Ago2 KO cells (13). In this study we have defined the locations on the HCV genome to which small RNA annealing induces HCV replication further defined the underlying mechanism of replication promotion by miR-122. We identified that small RNA annealing to nucleotides 1-3 and 15 - 44 promote HCV replication and that annealing to nucleotide 45 (SLIIa) and beyond do not (Figure 1 and 7). Further siRNAs binding to 15 nucleotides is required (Figures 6). We found that the ability of siRNAs to promote replication is related to their ability to induce the canonical 5’ UTR RNA structure, including formation of SLI, SLIIa, and SLIIb (Figure 5), (Supplementary Tables 1. and 2). This suggests that location specific binding of small RNAs promotes virus replication by favoring the formation of the correct 5’ UTR and IRES RNA structures.

We also determined that siRNA binding to other regions on HCV genome, including the HCV IRES does not stimulate virus replication. In addition, IRES annealing siRNAs neither promoted nor inhibited HCV replication (Figure 3C) and thus do not appear to disrupt IRES structure and function. Therefore, small RNA annealing induced RNA structure changes appear to be specific to the HCV 5’ terminal region and the HCV IRES structures may be too stable to be disrupted by small RNA annealing.

At least 7 more miR-122 binding sites were predicted in the HCV genome and were speculated to also affect the virus life cycle, (11, 40, 41, 43). However, none of the siRNAs that bound to the predicted miR-122 binding sites promoted replication (Figure 4). Our data therefore suggests that the two miR-122 binding sites on HCV 5’UTR are the only active binding sites that promote HCV replication. This data is in agreement with a recently published report in which mutation of the other miR-122 binding sites had no influence on HCV replication (40).

The minimum annealing required for efficient small RNA induced HCV replication promotion was 15 nucleotides. (Figure 6A and B). This data may explain why annealing of two copies of miR-122 is required for efficient HCV replication since annealing of one copy does not fulfill this requirement but annealing of two copies does (Figure 1). Further, the most efficient HCV promoting small RNA identified in our siRNA walk (si19-37) anneals to nucleotides comprising miR-122 binding seed site 1 (nucleotides 21-27) and the accessory miR-122 binding site 2 (nucleotides 29-31) and suggests that binding to miR-122 binding seed site 1 and accessory site 2 may be the minimum requirement for efficient small RNA dependent HCV replication. This data also supports a previous finding that miR-122 site 1 behaves similar to a conventional miRNA:target interaction where binding to a seed site is important (45), and that miR-122 binding site 2 has higher affinity owing to extended base pairing to the accessory site.

A previous report showed that miR-122 binding to the extreme 5’ terminus of HCV genome was required for efficient HCV replication (20). Our data indicates that small RNA annealing to the 5’ terminal region is not essential but enhances HCV replication promotion (Figure 7A and B). Our structure predictions also showed that siRNAs that interacted with 5’ terminal nucleotide induce the canonical IRES structure (Figure 7C). We therefore speculate that small RNA binding to terminal nucleotides on HCV genome may induce the canonical HCV structure better than those that do not. The previous report also showed that the 7 nucleotide 3’miR-122 overhang contributes to virus replication (20). However, our study indicates that a 2UU overhang was sufficient to promote HCV replication, and that generation of a miR-122-like 3’ overhang was not necessary and, in fact, was detrimental to HCV replication (Figure 7D and E).

All small RNAs that promote HCV replication were predicted to induce the canonical HCV IRES structure (Figure 2D and 5), (Supplementary Table 1). We therefore hypothesized that the ability of a small RNA to induce the IRES and thus stimulate translation will correlate with its ability to promote virus replication. Our data support this hypothesis and showed that siRNAs that promote replication efficiently also stimulated translation efficiently, and that translation stimulation correlated with virus replication levels (Figure 8C and D), (Supplementary Table 1). This suggests that translation stimulation as a key mechanism by which small RNA annealing promotes HCV replication. This data suggests that the miR-122 binding sites might be considered part of the IRES, or an IRES modulator.

Annealing of miR-122 to the HCV 5’ UTR stabilizes the viral genome, and this has been proposed as a mechanism by which miR-122 promotes HCV replication (13, 25). Further, it was speculated that the mechanism of protection is based on the generation of a double stranded 5’ terminus by miR-122 binding which protect it from cellular pyrophosphatases and exonuclease (Figure 2A) (20). We showed previously that si19-37 also stabilized HCV RNA suggesting that small RNAs may also promote virus life cycle by stabilizing the genome. However, si19-37 does not bind to the extreme 5’ terminus thus indicating that end annealing and the generation of an overhang is not required for genome stabilization. Further, we show that siRNAs annealing stabilizes the HCV genome regardless of whether they promote replication (Figure 9). Therefore, genome stabilization is not sufficient alone to promote virus replication, but functions to enhance replication induced by translation stimulation.

**Figure 9:**
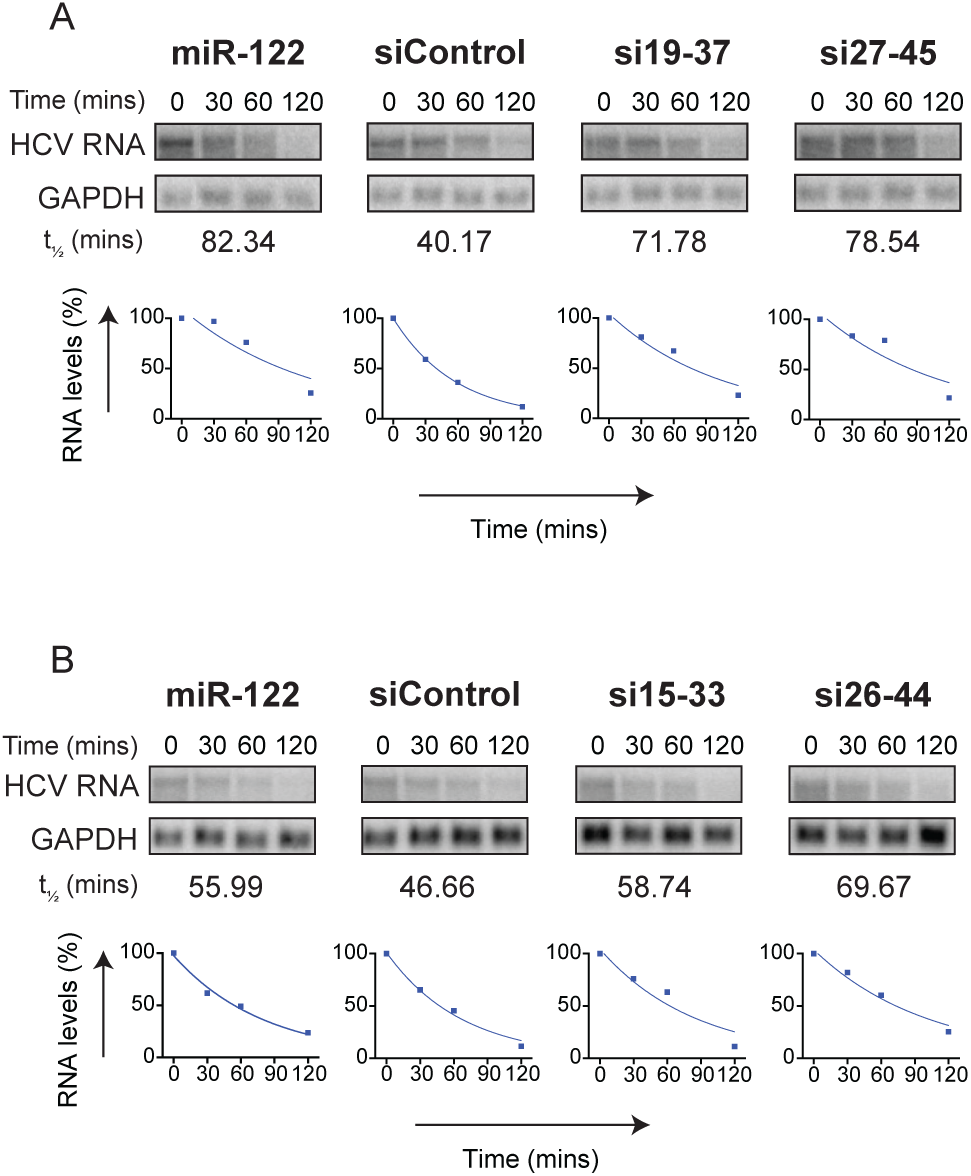
Genome stabilization by small RNAs that do and do not promote HCV replication. (A and B) Northern blot analyses of HCV genomic RNA quantities during stability assays. Assays are shown for HCV RNA with annealing of miR-122, two different siRNAs, or siControl. Bands were quantified using Image-Studio Lite and were plotted as a one phase decay curve. These data are representative of 2 independent experiments performed with each of the 4 siRNAs along with control small RNAs (miR122 and sicontrol). Decay curves for each sample (miR-122/siControl/19-37/si27-45/si15-33/si26-44) were generated independently and half-lives obtained from these decay curves are mentioned. Full uncropped blots are shown in Supplement Figure 2A. Decay curves and half-lives were calculated by Graphpad Prism.

Based on our findings we present a model in which position-specific annealing of small RNAs induces the formation of the viral IRES RNA structures and stimulates virus translation. In addition, position-independent small RNA annealing stabilizes the viral genome but alone is insufficient to promote the virus life cycle (Figure 10). However, it is still unknown whether HCV genome structure changes induced by small RNA binding are fixed throughout the virus lifecycle or whether they are dynamic and differ during specific events in the virus lifecycle such as replication and virion assembly. In addition, the roles of miRNA associated proteins like Ago and Ago complexes in HCV promotion by miR-122 remain to be clarified.

**Figure 10:**
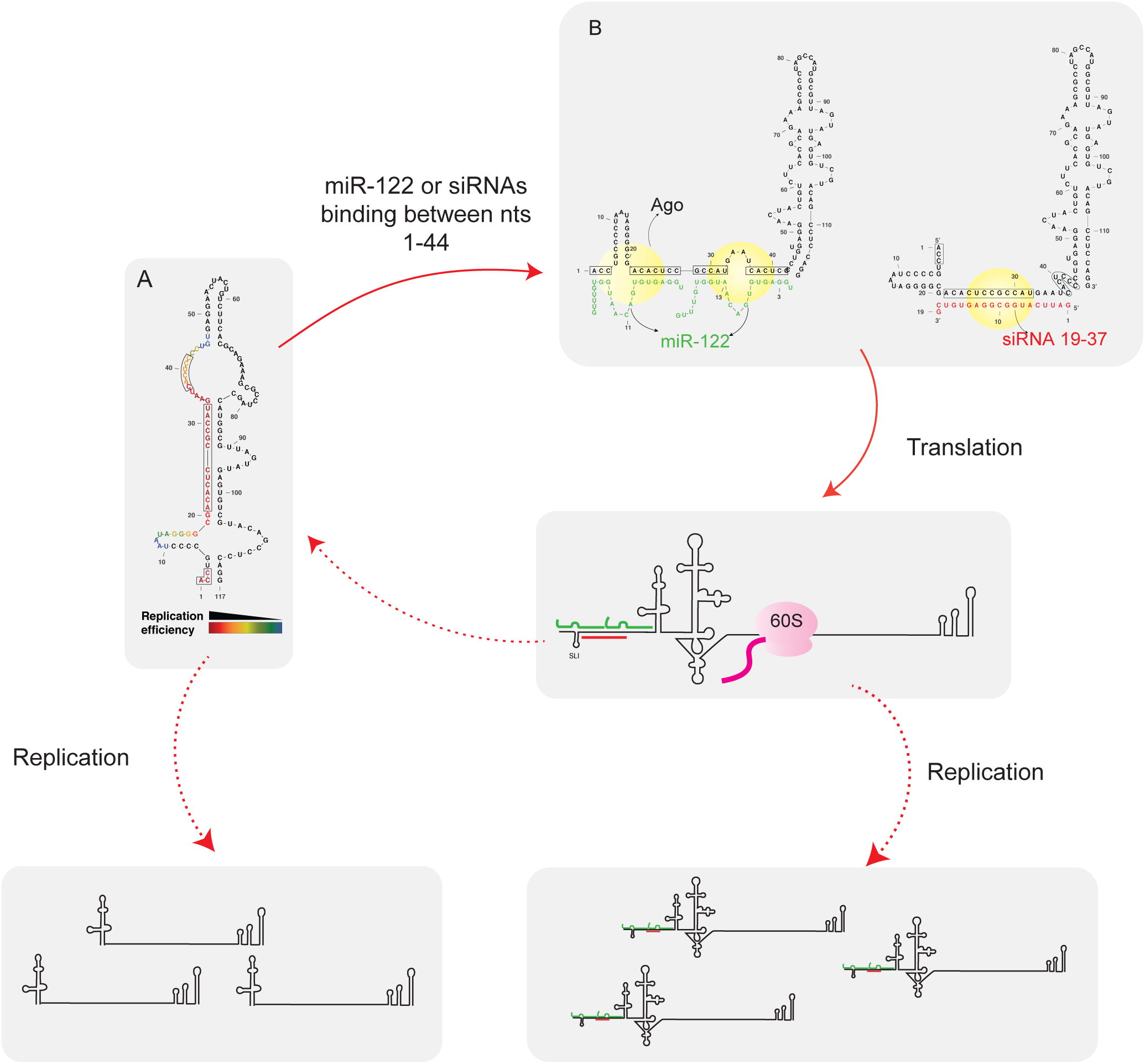
Summary model figure: We have developed a model for the mechanism of miR-122 promotion of HCV. In our model, the (A). HCV 5’ UTR RNA forms a non-canonical structure (colours of nucleotides represent small RNA replication efficiency when bound to those nucleotides) in the absence of small RNA annealing but (B). the canonical 5’ UTR and IRES structure when bound with 2 copies of miR-122 or with siRNAs. (A and B). Boxed nucleotides represent miR122 binding sites. Small RNA induction or stabilization of the canonical IRES structure promotes virus translation leading to enhanced virus replication. Small RNA annealing also stabilizes the viral genome, but genome stabilization alone is not sufficient to promote the HCV lifecycle. We propose that small RNAs anneal associate with host Argonaute proteins and together they are responsible for RNA structure changes and viral genome stabilization. It is unknown if genome amplification is affected by miR-122 or whether replication initiation is regulated by the canonical or non-canonical

## ACKNOWLEDGEMENTS

We would like to acknowledge Charlie Rice (The Rockefeller University) for providing the pJ6/JFH-1(p7Rluc2A), DROSHA knockout, and Huh-7.5 cells. We also thank Matthew Evans for miR-122 knockout cells.

## Authors’ contribution

R.K., S.G., J.Q.K. and J.A.W. designed and performed the experiments, and analyzed the data; R.K. and J.A.W. wrote the manuscript.

---

**Supplement Figure 1:**
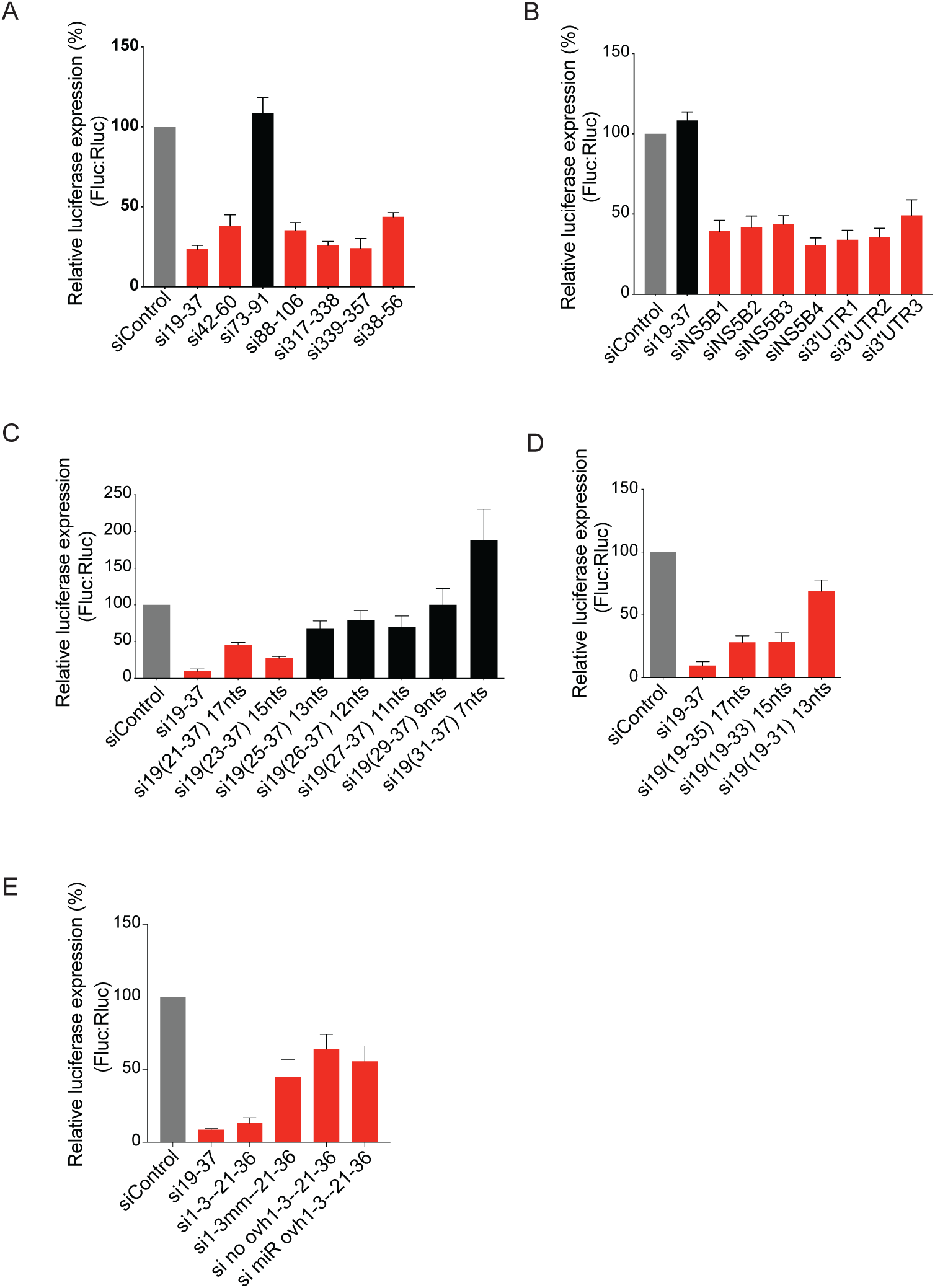
SiRNA suppression assays for A.) siRNAs that bind within the HCV IRES B.) siRNAs that bind within NS5B and the 3’UTR C and D.) si19-37 analogues E.) siRNAs tested for terminal binding and overhang. Red bars indicate siRNAs that suppress translation, black bars indicate siRNAs that do not suppress translation.

**Supplement Figure 2:**
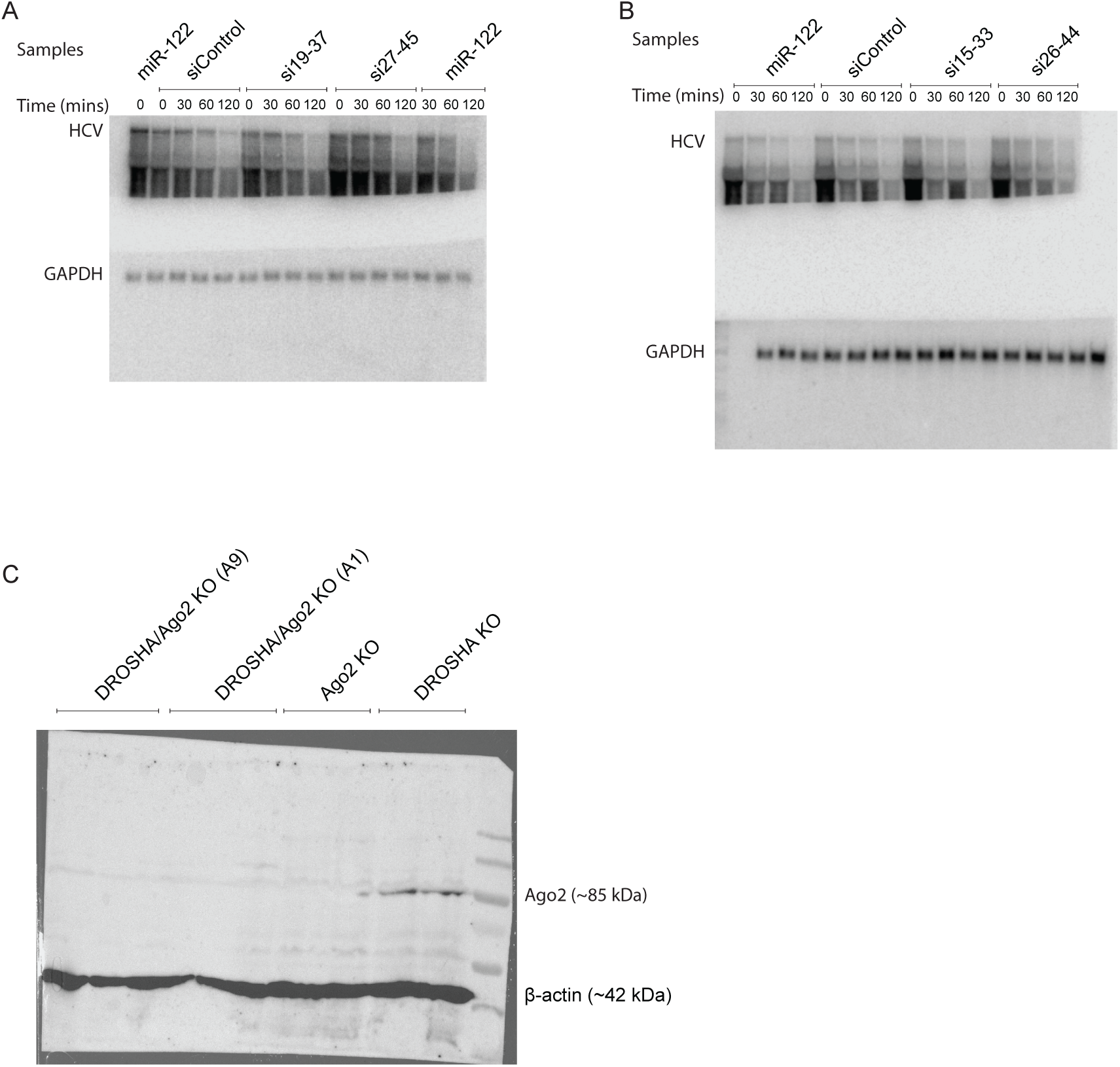
A.) and B.) Northern blot membranes for experiments describes in Stabilization assay. C.) Western blot described to assess expression of Ago2 protein in DROSHA/Ago2 KO cells. DROSHA/Ago2 cell clone A9 was used for our studies.

**Supplementary Table 1:**
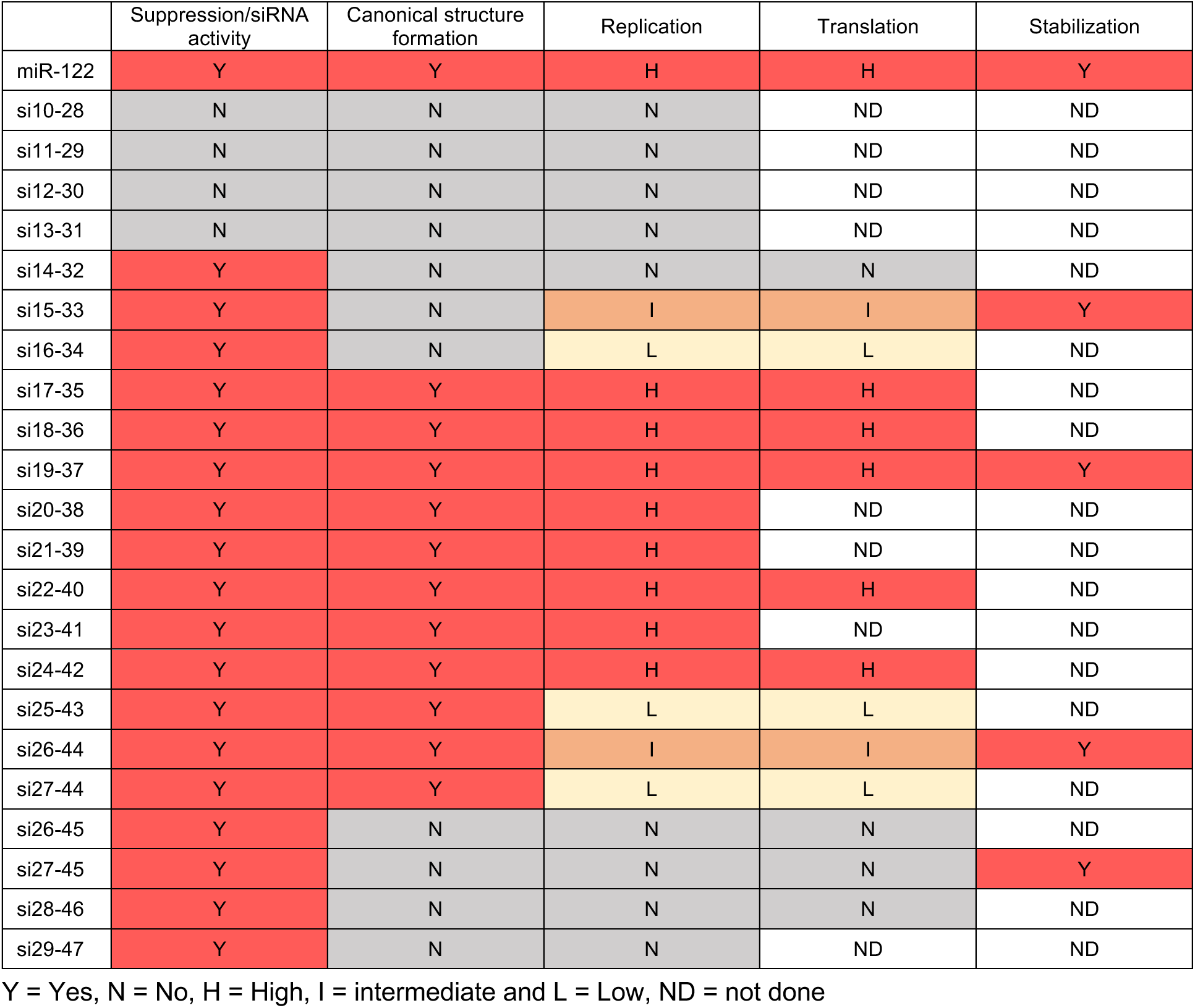

**Supplementary Table 2:**
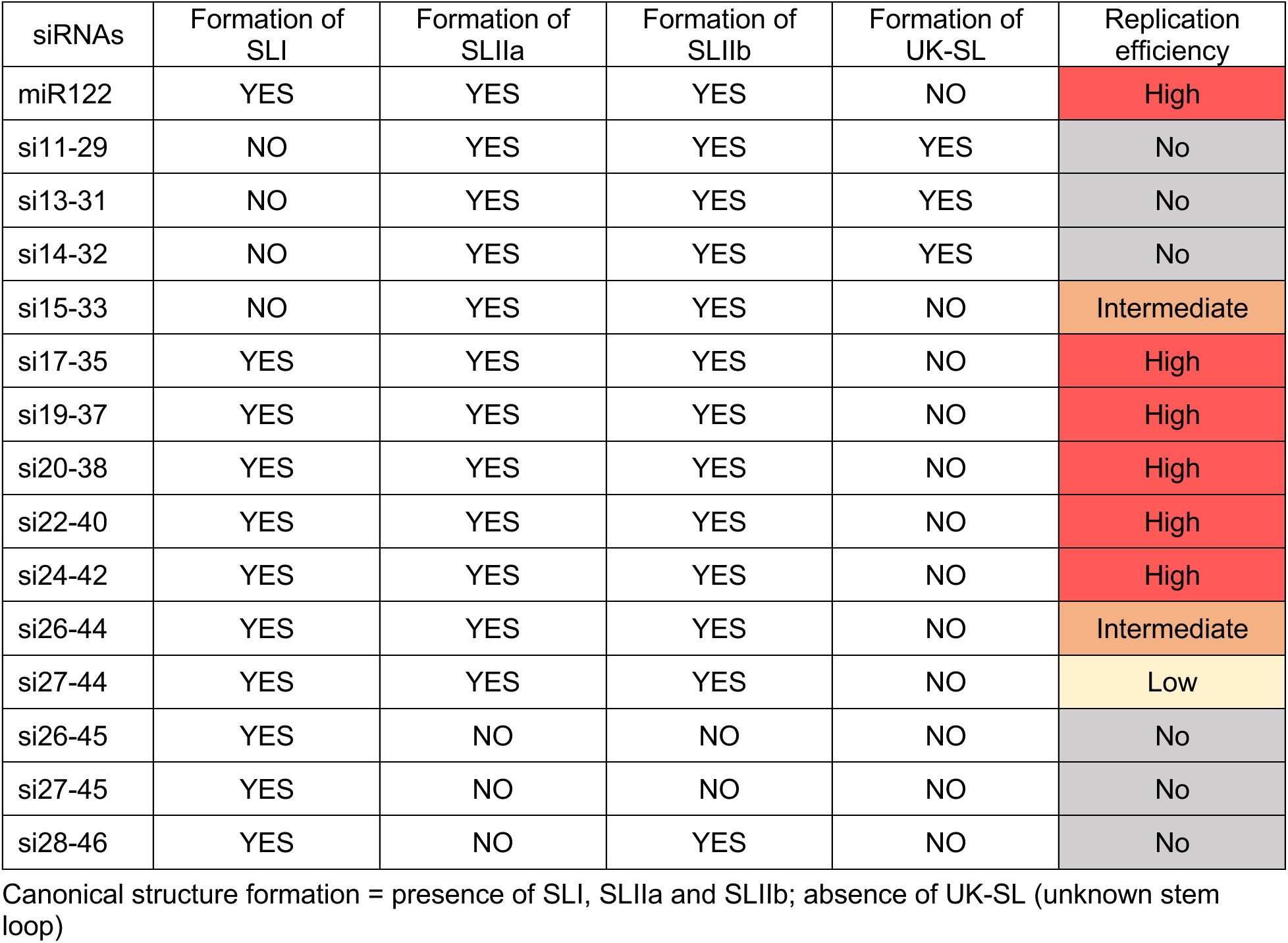

**Supplementary Table 3:**
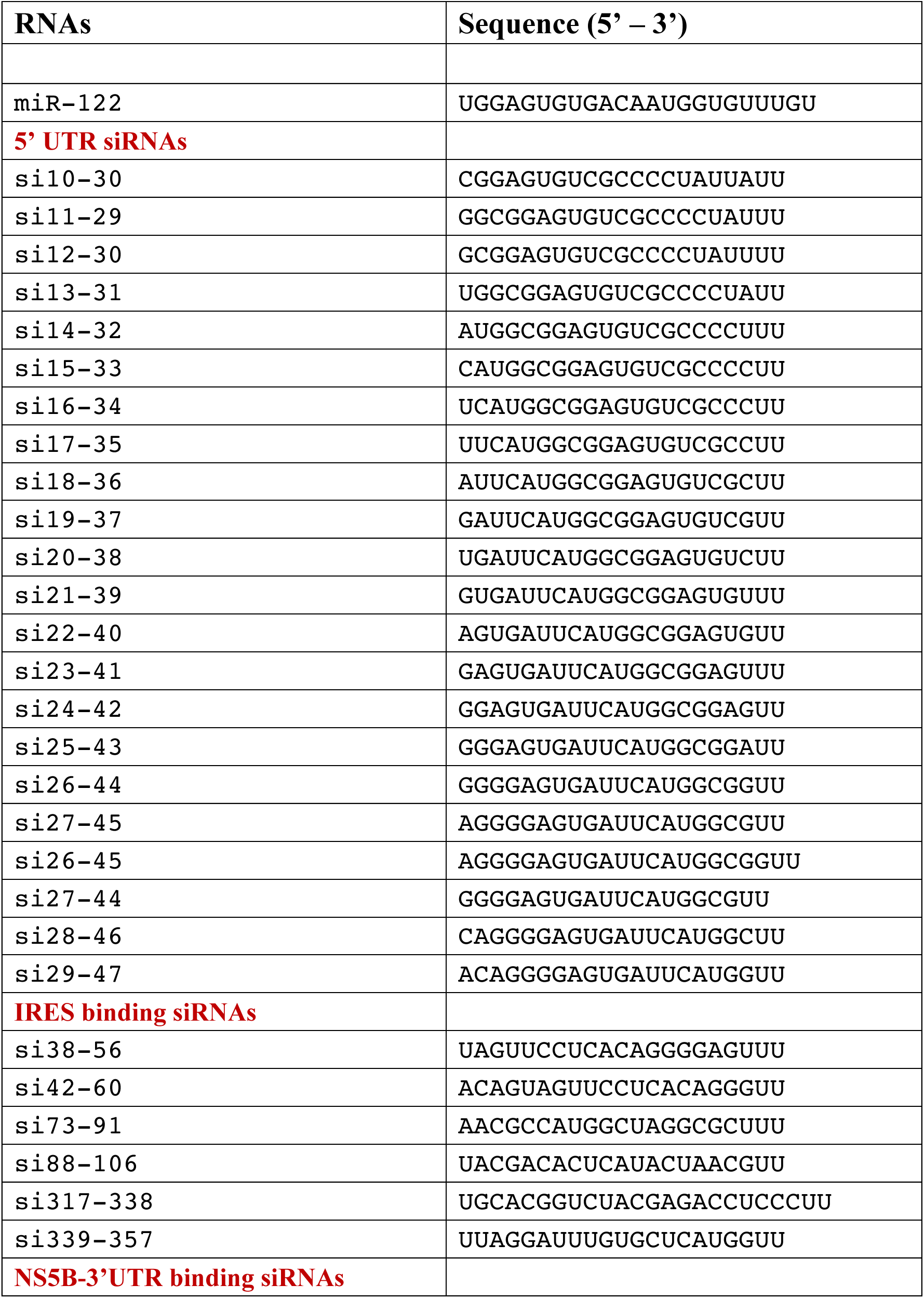
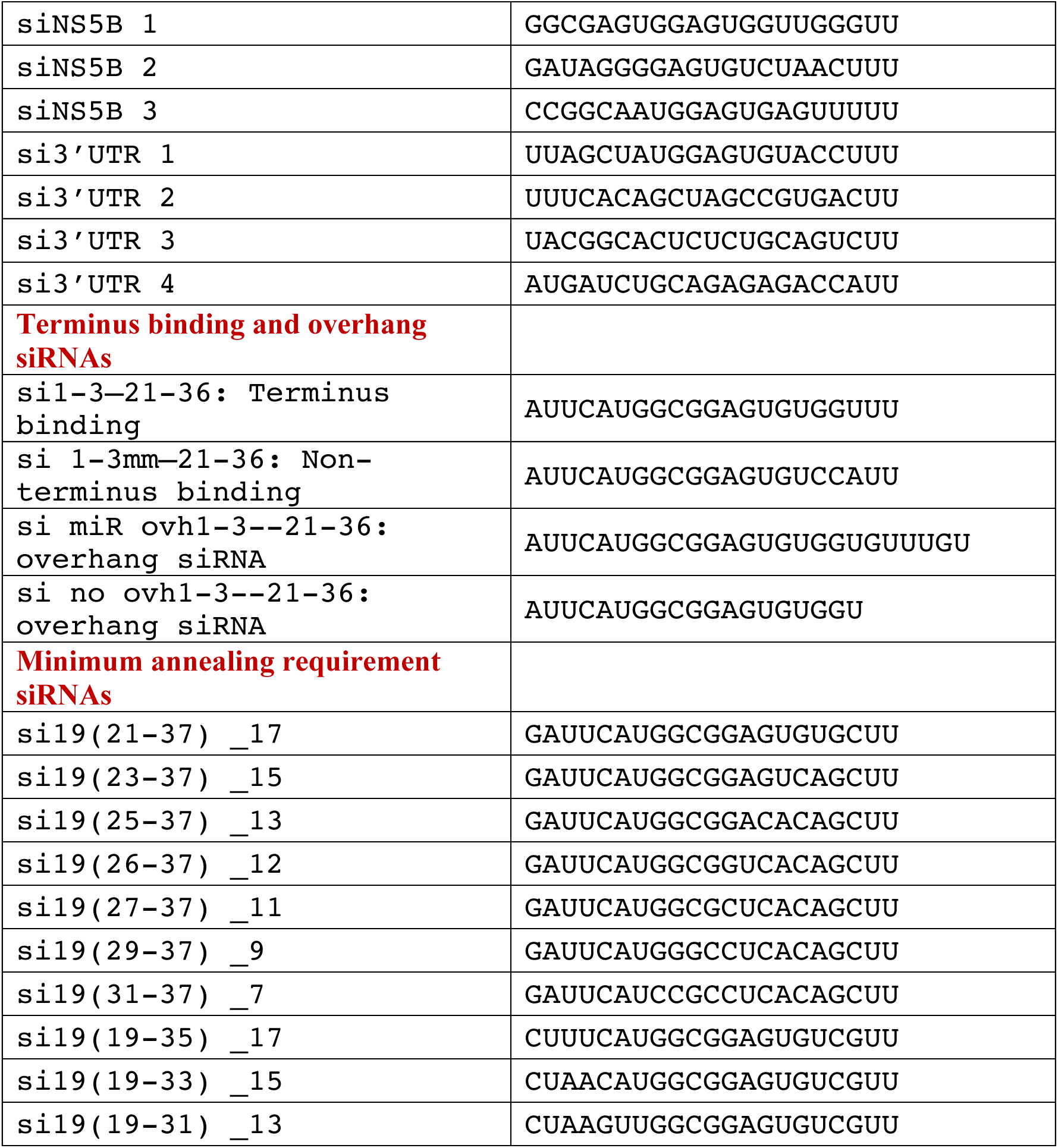
small RNAs used in this study

