## Supplemental Data for "Location specific small RNA annealing to the HCV 5’ UTR promotes Hepatitis C Virus replication by favoring IRES formation and stimulating virus translation"

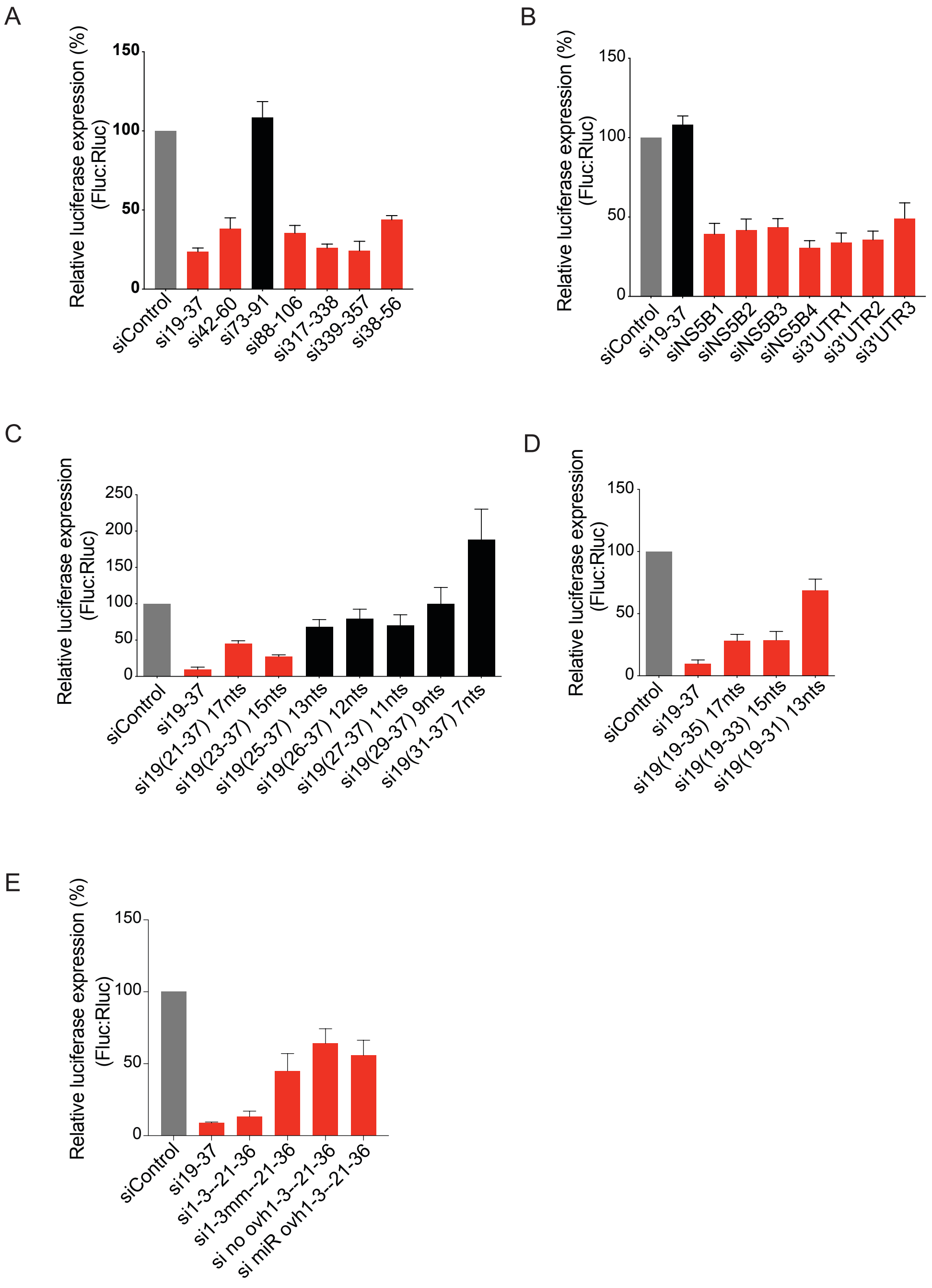

**Supplement Figure 1:** SiRNA suppression assays for A.) siRNAs that bind within the HCV IRES B.) siRNAs that bind within NS5B and the 3'UTR C and D.) si19-37 analogues E.) siRNAs tested for terminal binding and overhang. Red bars indicate siRNAs that suppress translation, black bars indicate siRNAs that do not suppress translation.

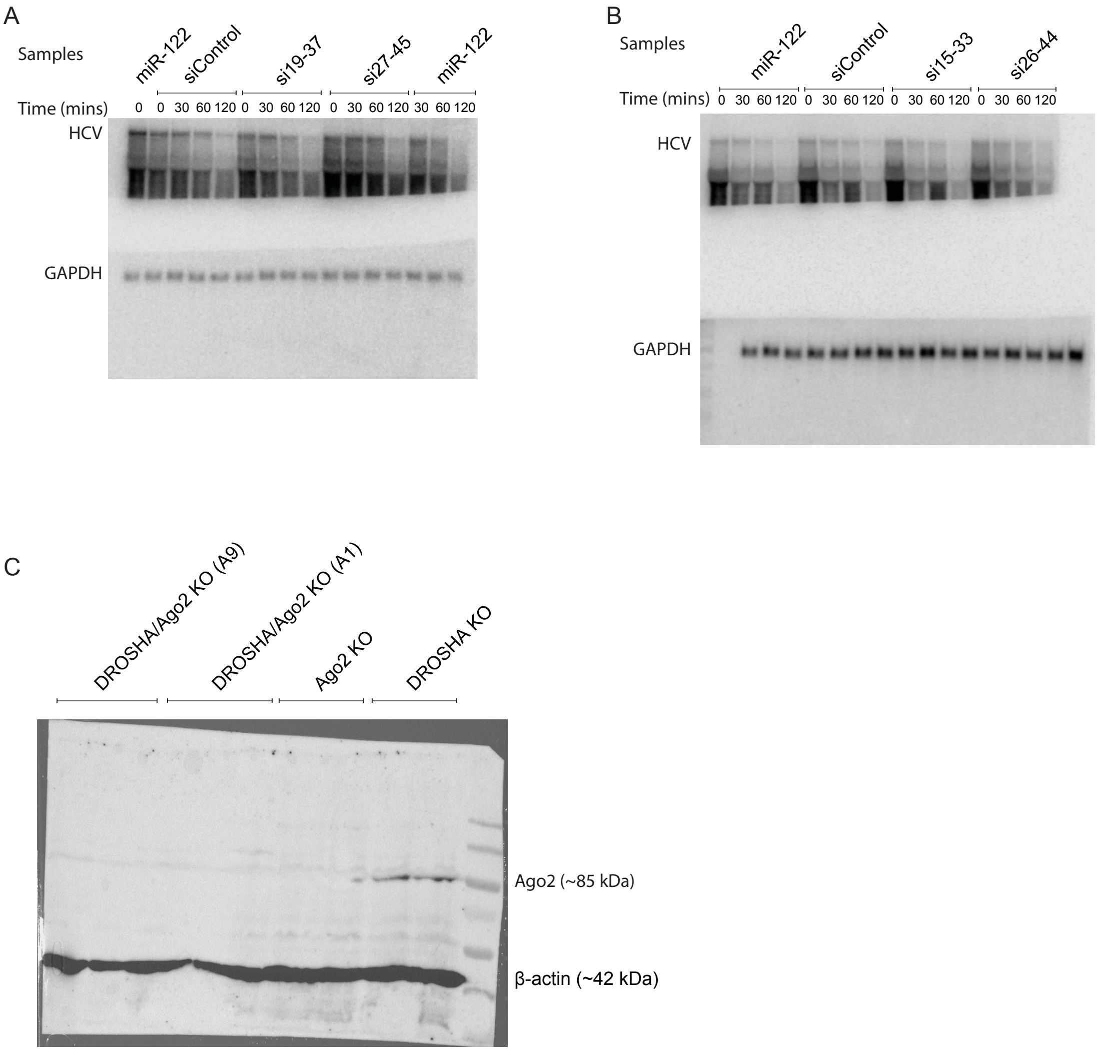

**Supplement Figure 2:** A.) and B.) Northern blot membranes for experiments describes in Stabilization assay. C.) Western blot described to assess expression of Ago2 protein in DROSHA/Ago2 KO cells. DROSHA/Ago2 cell clone A9 was used for our studies.

### Supplementary Table 1:

| | Suppression/siRNA activity | Canonical structure formation | Replication | Translation | Stabilization |
|---|---|---|---|---|---|
| miR-122 | Y | Y | H | H | Y |
| si10-28 | N | N | N | ND | ND |
| si11-29 | N | N | N | ND | ND |
| si12-30 | N | N | N | ND | ND |
| si13-31 | N | N | N | ND | ND |
| si14-32 | Y | N | N | N | ND |
| si15-33 | Y | N | I | I | Y |
| si16-34 | Y | N | L | L | ND |
| si17-35 | Y | Y | H | H | ND |
| si18-36 | Y | Y | H | H | ND |
| si19-37 | Y | Y | H | H | Y |
| si20-38 | Y | Y | H | ND | ND |
| si21-39 | Y | Y | H | ND | ND |
| si22-40 | Y | Y | H | H | ND |
| si23-41 | Y | Y | H | ND | ND |
| si24-42 | Y | Y | H | H | ND |
| si25-43 | Y | Y | L | L | ND |
| si26-44 | Y | Y | I | I | Y |
| si27-44 | Y | Y | L | L | ND |
| si26-45 | Y | N | N | N | ND |
| si27-45 | Y | N | N | N | Y |
| si28-46 | Y | N | N | N | ND |
| si29-47 | Y | N | N | ND | ND |

Y = Yes, N = No, H = High, I = intermediate and L = Low, ND = not done

**Supplementary Table 2:**

| siRNAs | Formation of SLI | Formation of SLIIa | Formation of SLIIb | Formation of UK-SL | Replication efficiency |
|---|---|---|---|---|---|
| miR122 | YES | YES | YES | NO | High |
| si11-29 | NO | YES | YES | YES | No |
| si13-31 | NO | YES | YES | YES | No |
| si14-32 | NO | YES | YES | YES | No |
| si15-33 | NO | YES | YES | NO | Intermediate |
| si17-35 | YES | YES | YES | NO | High |
| si19-37 | YES | YES | YES | NO | High |
| si20-38 | YES | YES | YES | NO | High |
| si22-40 | YES | YES | YES | NO | High |
| si24-42 | YES | YES | YES | NO | High |
| si26-44 | YES | YES | YES | NO | Intermediate |
| si27-44 | YES | YES | YES | NO | Low |
| si26-45 | YES | NO | NO | NO | No |
| si27-45 | YES | NO | NO | NO | No |
| si28-46 | YES | NO | YES | NO | No |

Canonical structure formation = presence of SLI, SLIIa and SLIIb; absence of UK-SL (unknown stem loop)

**Supplementary Table 3: small RNAs used in this study**

| RNAs | Sequence (5' – 3') |
|---|---|
| miR-122 | UGGAGUGUGACAAUGGUGUUUGU |
| **5' UTR siRNAs** | |
| si10-30 | CGGAGUGUCGCCCCUAUUAUU |
| si11-29 | GGCGGAGUGUCGCCCCUAUUU |
| si12-30 | GCGGAGUGUCGCCCCUAUUUU |
| si13-31 | UGGCGGAGUGUCGCCCCUAUU |
| si14-32 | AUGGCGGAGUGUCGCCCCUUU |
| si15-33 | CAUGGCGGAGUGUCGCCCCUU |
| si16-34 | UCAUGGCGGAGUGUCGCCCUU |
| si17-35 | UUCAUGGCGGAGUGUCGCCUU |
| si18-36 | AUUCAUGGCGGAGUGUCGCUU |
| si19-37 | GAUUCAUGGCGGAGUGUCGUU |
| si20-38 | UGAUUCAUGGCGGAGUGUCUU |
| si21-39 | GUGAUUCAUGGCGGAGUGUUU |
| si22-40 | AGUGAUUCAUGGCGGAGUGUU |
| si23-41 | GAGUGAUUCAUGGCGGAGUUU |
| si24-42 | GGAGUGAUUCAUGGCGGAGUU |
| si25-43 | GGGAGUGAUUCAUGGCGGAUU |
| si26-44 | GGGGAGUGAUUCAUGGCGGUU |
| si27-45 | AGGGGAGUGAUUCAUGGCGUU |
| si26-45 | AGGGGAGUGAUUCAUGGCGGUU |
| si27-44 | GGGGAGUGAUUCAUGGCGUU |
| si28-46 | CAGGGGAGUGAUUCAUGGCUU |
| si29-47 | ACAGGGGAGUGAUUCAUGGUU |
| **IRES binding siRNAs** | |
| si38-56 | UAGUUCCUCACAGGGGAGUUU |
| si42-60 | ACAGUAGUUCCUCACAGGGUU |
| si73-91 | AACGCCAUGGCUAGGCGCUUU |
| si88-106 | UACGACACUCAUACUAACGUU |
| si317-338 | UGCACGGUCUACGAGACCUCCCUU |
| si339-357 | UUAGGAUUUGUGCUCAUGGUU |
| **NS5B-3'UTR binding siRNAs** | |

| | |
|---|---|
| siNS5B 1 | GGCGAGUGGAGUGGUUGGGUU |
| siNS5B 2 | GAUAGGGGAGUGUCUAACUUU |
| siNS5B 3 | CCGGCAAUGGAGUGAGUUUUU |
| si3'UTR 1 | UUAGCUAUGGAGUGUACCUUU |
| si3'UTR 2 | UUUCACAGCUAGCCGUGACUU |
| si3'UTR 3 | UACGGCACUCUCUGCAGUCUU |
| si3'UTR 4 | AUGAUCUGCAGAGAGACCAUU |
| **Terminus binding and overhang siRNAs** | |
| si1-3—21-36: Terminus binding | AUUCAUGGCGGAGUGUGGUUU |
| si 1-3mm—21-36: Non-terminus binding | AUUCAUGGCGGAGUGUCCAUU |
| si miR ovh1-3--21-36: overhang siRNA | AUUCAUGGCGGAGUGUGGUGUUUGU |
| si no ovh1-3--21-36: overhang siRNA | AUUCAUGGCGGAGUGUGGU |
| **Minimum annealing requirement siRNAs** | |
| si19(21-37) _17 | GAUUCAUGGCGGAGUGUGCUU |
| si19(23-37) _15 | GAUUCAUGGCGGAGUCAGCUU |
| si19(25-37) _13 | GAUUCAUGGCGGACACAGCUU |
| si19(26-37) _12 | GAUUCAUGGCGGUCACAGCUU |
| si19(27-37) _11 | GAUUCAUGGCGCUCACAGCUU |
| si19(29-37) _9 | GAUUCAUGGGCCUCACAGCUU |
| si19(31-37) _7 | GAUUCAUCCGCCUCACAGCUU |
| si19(19-35) _17 | CUUUCAUGGCGGAGUGUCGUU |
| si19(19-33) _15 | CUAACAUGGCGGAGUGUCGUU |
| si19(19-31) _13 | CUAAGUUGGCGGAGUGUCGUU |
